# Socio-environmental and measurement factors drive spatial variation in influenza-like illness

**DOI:** 10.1101/112680

**Authors:** Elizabeth C. Lee, Ali Arab, Sandra Goldlust, Cécile Viboud, Shweta Bansal

## Abstract

The mechanisms hypothesized to drive spatial heterogeneity in reported influenza activity include: environmental factors, contact patterns, population age structure, and socioeconomic factors linked to healthcare access and quality of life. Harnessing the large volume and high specificity of diagnosis codes in medical claims data for influenza seasons from 2002-2009, we estimate the importance of socio-environmental determinants and measurement-related factors on observed variation in influenza-like illness (ILI) across United States counties. We found that South Atlantic states tended to have higher ILI seasonal intensity, and a combination of transmission, environmental, influenza subtype, socioeconomic and measurement factors explained the variation in seasonal intensity across our study period. Moreover, our models suggest that sentinel surveillance systems should have fixed report locations across years for the most robust inference and prediction, and high volumes of data can offset measurement biases in opportunistic data samples.

## Introduction

Seasonal influenza represents an important public health burden worldwide, and even within a single year, there is substantial variation in disease burden across populations (*Moorthy et al., 2012*; *Lee et al., 2015*). Many studies have examined the drivers and patterns influenza seasonality (*Lofgren et al., 2007; Tamerius et al., 2011*), while others have focused on the large-scale spatial patterns in influenza epidemic timing, suggesting for instance, spread from West to East across North America due to a combination of local contact patterns and global travel patterns (*Wenger and Naumova, 2010; Schanzer et al., 2011b;Grais et al., 2003; Brownstein et al., 2006*). While there are numerous studies explaining spatial variation in seasonal influenza transmission and disease burden, most studies focus on very aggregated or very local study areas (e.g., country-level or one school district, respectively) compare only one or two hypotheses in isolation.

Among these, humidity and temperature have each been associated with seasonal flu onset, seasonal fluctuations, and heightened morbidity and mortality in epidemiological contexts *(Shaman et al., 2010;Yu et al., 2013;Barreca and Shimshack, 2012;Deyle et al., 2016),* and lower humidity and colder temperatures may increase influenza virus transmission and survival (*Lowen et al., 2007; Shaman and Kohn, 2009*) Chronic illnesses such as asthma, exacerbated by air pollution, elevated the risk for severe symptoms of pandemic H1N1 (*Van Kerkhove et al., 2011*) Empirical evidence supports the occurrence of both aerosol and droplet transmission of influenza virus (*Killingley and Nguyen-Van-Tam, 2013*) and these transmission modes suggest that influenza seasons may follow both density-dependent and frequency-dependent disease dynamics (per capita contact rates between susceptible and infectious individuals do and do not change with population density, respectively). The high connectivity of school-aged children in contact surveys (*Mossong et al., 2008*; *Kucharski et al., 2014*) has led to hypotheses that children drive local transmission and adults seed new infections across longer distances (*Viboud et al., 2006; Apolloni et al., 2013*) which may manifest in shifted epidemic timings across age groups (*Lemaitre and Carrat, 2010;Peters et al., 2014;Schanzer et al., 2010;Wallinga et al., 2006; Timpka et al., 2012*) Immune landscapes vary across locations; epidemic outcomes in one season may trickle down to subsequent years through differences in cross-protective immunity, and high flu vaccination coverage may reduce morbidity and incidence of severe clinical outcomes (*Kostova et al., 2013*) Finally, flu type and subtype circulation may also drive spatial heterogeneity; A/H3-dominant flu seasons are associated with greater morbidity and mortality and an older patient age distribution than A/H1 season (*Frank et al., 1985;Simonsen et al., 1997;Khiabanian et al., 2009; Peters et al., 2014*) while influenza B is thought to circulate predominantly and earlier among children (*Peters et al., 2014;Hayward et al., 2014; Beauté et al., 2015*)

Beyond socio-environmental mechanisms, we must consider the possibility that the measurement of influenza disease burden plays a significant role in driving the observed spatial heterogeneity. While poverty and other social determinants are thought to increase risk for influenza morbidity, hospitalization, and mortality (*Lowcock et al., 2012;Kumar et al., 2015;Hadler et al., 2016;Charland et al., 2011; Grantz et al., 2016*) these observations are often confounded by care-seeking behavior, the likelihood that sick individuals will seek treatment from a health care provider. Roughly 43% of adults and 60% of elderly seek care for influenza-like illness (ILI) in the United States, as many cases are too mild to warrant a visit to the doctor (*Biggerstaff et al., 2014b*). In addition to differences in personal choice, limited access to health care and health insurance also delay or reduce care-seeking behavior, further generating biases in reported case severity or patient numbers among physician-based surveillance systems (*Biggerstaff et al., 2014b*)

In this study, we examine the transmission, environmental, influenza-specific, and socioeconomic mechanisms and measurement processes underlying the spatial variation in reported influenza-like illness across counties in the United States. Leveraging highly resolved medical claims data, we identified important drivers of spatial heterogeneity in the magnitude and duration of flu seasons from 2002 to 2009 in a large-scale ecological analysis. We then used our Bayesian modeling framework in new applications to probe the robustness of this ecological inference with limited data availability and to assess the predictive ability of our model in a more recent flu season. Our results highlight the relative contributions of surveillance data collection and socio-environmental processes to disease reporting, and highlight the importance of considering measurement biases when using surveillance data for epidemiological inference and prediction.

## Results

We examined the socio-environmental and measurement-related drivers of spatial heterogeneity in influenza disease burden across U.S. counties for flu seasons from 2002-2003 through 2008-2009 using a hierarchical Bayesian modeling approach. Using medical claims data representing 2.5 billion visits from upwards of 120,000 health care providers each year, our study considered six disease burden response variables: two measures of influenza disease burden (relative risk of seasonal intensity, which is a proxy for attack rate, and epidemic duration in number of weeks) in three populations (total population, children 5-19 years old, and adults 20-69 years old) with multi-season and single season model structures. There were 13 county-level, 2 state-level and 4 HHS region-level predictors in the final model *Table 1;* all predictors were the same across response variables except care-seeking behavior, which was specific to the age group in the response. The seasonal intensity model fit the data well and the Pearson’s cross-correlation coefficient between the log seasonal intensity and log prediction was *R* = 0.87 (*Figure 1*). Results reported in the following sections are from the multi-season total population seasonal intensity model unless otherwise noted.

**Table 1.** Final model predictors and hypotheses.

| Factor | Index | Plot Label | Spatial Scale | Hypothesized Effect |
| --- | --- | --- | --- | --- |
| **Environmental factors** |  |  |  |  |
| Flu transmission | Specific humidity | humidity | county | – |
| Respiratory disease risk | Fine particular matter | pollution | county | + |
| **Transmission mechanisms** |  |  |  |  |
| Density-dependent | Population density | popDensity | county | + |
| Frequency-dependent | Average household size | householdSize | county | + |
| **Diffusion mechanisms** |  |  |  |  |
| Local spread | % child population | child | county | + |
| Importation risk | % adult population | adult | county | + |
| **Immunity** |  |  |  |  |
| Vaccine-acquired | Toddler vacc. coverage | toddlerVacc | state | – |
|  | Elderly vacc. coverage | elderlyVacc | state | – |
| Prior exposure | Population protected due to prior season exposure | priorImmunity | county | – |
| **Influenza circulation** |  |  |  |  |
| Dominant A subtype | % H3 subtype among flu type A samples | fluH3 | HHS region | + |
| B circulation | % B type among positive flu samples | fluB | HHS region | + |
| H3 has older age distribution | adult population x dominant A subtype | adult-fluH3 | HHS region | + |
| B circulates primarily in children | child population x B circulation | child-fluB | HHS region | + |
| **Socioeconomic factors and access to care** |  |  |  |  |
| Health care availability | Hospitals per capita | hospAccess | county | + |
| Social deprivation | % single-person households | onePersonHH | county | + |
| Material deprivation | % in poverty | poverty | county | + |
| Claims-reporting population | % with health insurance | insured | county | + |
| **Measurement factors** |  |  |  |  |
| Claims database coverage | % physicians reporting to claims database | claimsCoverage | county | + |
| Care-seeking behavior in claims database | All visits per capita reported in database | careseeking | county | + |

**Figure 1.**
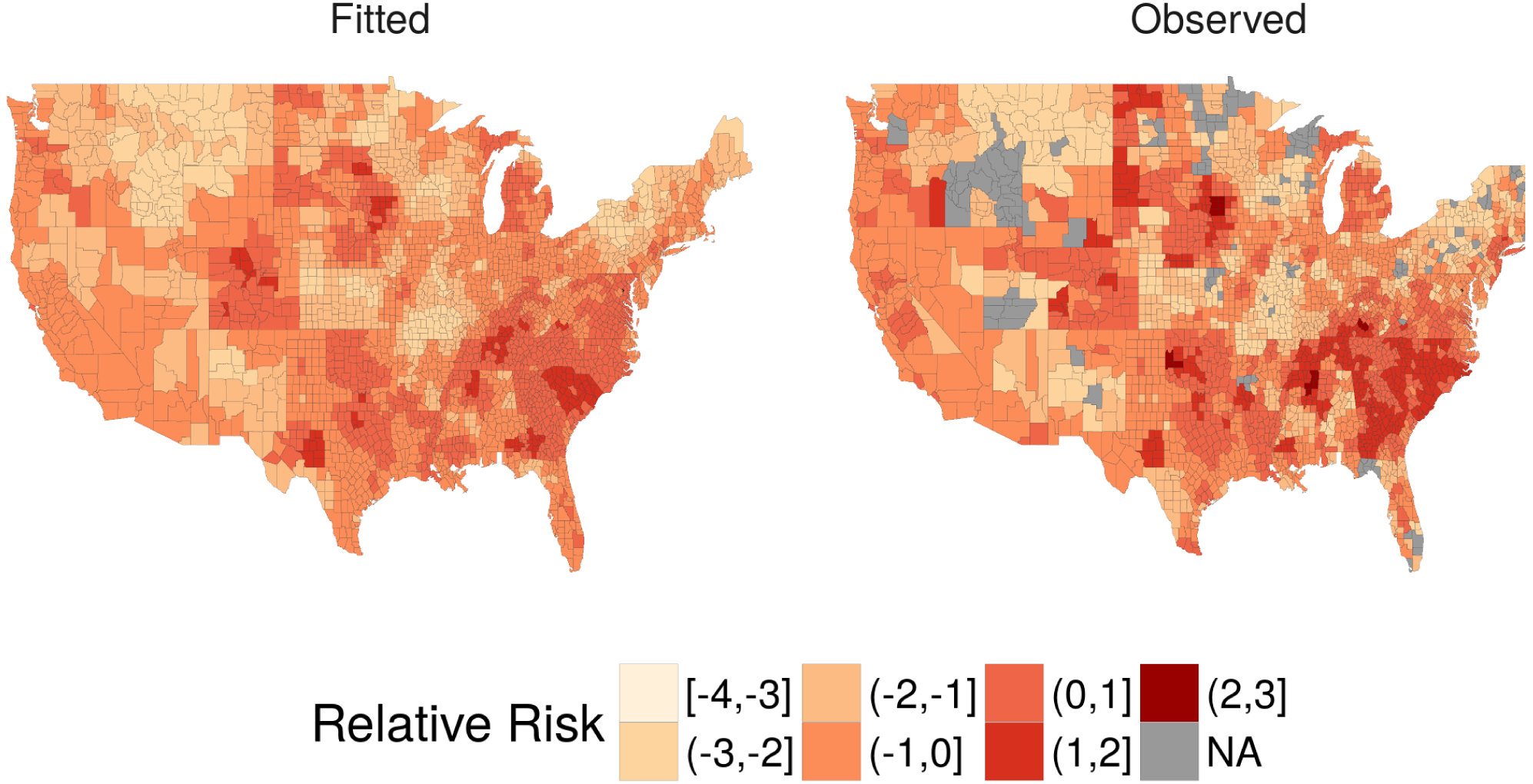
Continental U.S. county map for fitted and observed relative risk of seasonal intensity for an example flu season (2006-2007). **Figure 1-Figure supplement 1.** Continental U.S. county maps for fitted (left) and observed (right) relative risk of seasonal intensity for remaining influenza seasons.

### Temporal and spatial patterns of influenza-like illness

Group (random) effects were used to identify consistent spatial or temporal patterns across locations and study years. We found that the 2004-2005 flu season had greater seasonal intensity, while 2008-2009 had relatively low seasonal intensity (*Figure 2*). For the seasonal intensity model, no single region had a significant group effect, although several South Atlantic states like Georgia, Maryland, North Carolina, South Carolina, and Virginia had relatively greater risk than other states across the study period, while several Plains and Rocky Mountain states like Kansas, Minnesota, Missouri, Montana, and Utah had relatively lower risk.

**Figure 2.**
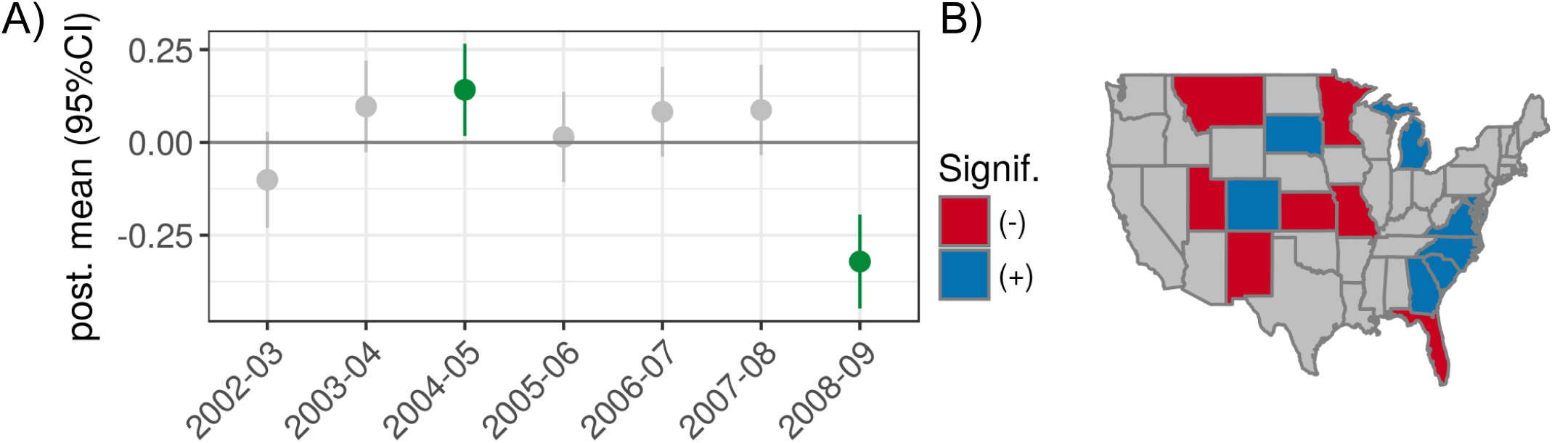
Temporal and spatial group effects for total population seasonal intensity. A) 95% credible intervals for group (random) effects by influenza season. B) Continental U.S. maps highlighting states with significantly greater or lower seasonal intensity across the study period.

### Drivers of seasonal intensity

Several socio-environmental drivers of seasonal intensity risk were identified in the multi-season model (*Figure 3*). Total seasonal intensity had positive associations with the adult-flu H3 and child-flu B interaction terms, estimated average household size, and a proxy for prior immunity. There were negative associations with adult and child population proportions, average flu season specific humidity, proportion of the population in poverty, proportion of single person households, and infant vaccination coverage.

**Figure 3.**
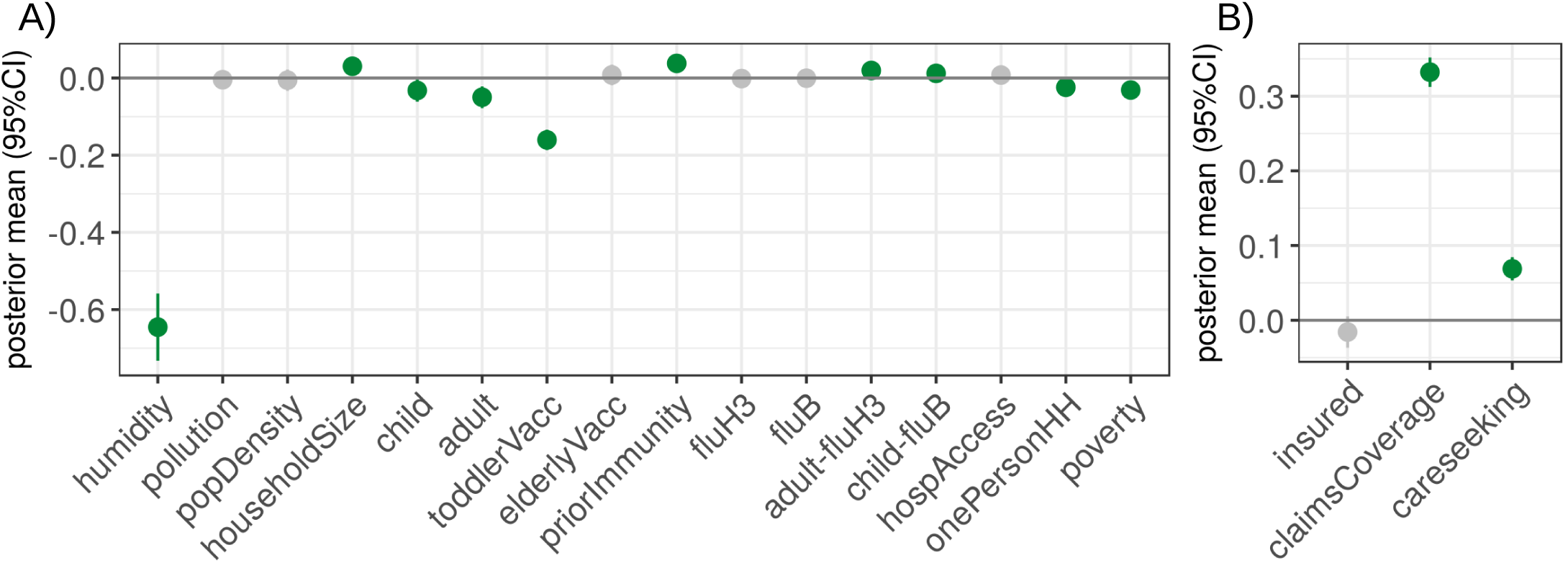
For the total population multi-season seasonal intensity models, these are the 95% credible intervals for the posterior distributions of the A) socio-environmental coefficients and B) measurement-related coefficients. Distributions indicated in green were statistically significant. **Figure 3-Figure supplement 1.** For the total population single-season seasonal intensity models, these are the 95% credible intervals for the posterior distributions of the socio-environmental coefficients. **Figure 3-Figure supplement 2.** For the total population single-season seasonal intensity models, these are the 95% credible intervals for the posterior distributions of the measurement coefficients.

We found that careseeking behavior and claims database coverage had strong positive associations with seasonal intensity (*Figure 3*). In considering the single-season models, the positive effect of claims database coverage on seasonal intensity appeared to decline in magnitude over time (*Figure 3* supplement). This corresponded with an increase in claims database coverage over time (*Appendix 5*).

### Drivers of age-specific seasonal intensity

Children and adults comprise the largest components of the U.S. population, and many studies have considered shifts in epidemic timing and immunity due to differences in contact patterns, shifting risk between children and adults over time, interactions between influenza types/subtypes by age, and differences in vaccine effectiveness by age group (*Bansal et al., 2010;Lee et al., 2015;Ewing et al., 2016; Schanzer et al., 2011a;Gostic et al., 2016; Khiabanian et al., 2009*). Considering the potential to elucidate age-specific transmission mechanisms and improve targeting of public health interventions, we used the multi-season model to examine drivers of seasonal intensity in the child and adult populations. Full model results are reported in *Appendix 2*, and for both age groups, predicted value means appeared to be systematically over-estimated relative to the observed relative risk of seasonal intensity. The Pearson’s cross-correlation coefficient between the log observation and log predicted mean was ***R*** = 0.89 and ***R*** = 0.90 for the child and adult seasonal intensity models, respectively.

Children had greater intensity in the 2003-2004 flu season and lower intensity in the 2002-2003 and 2008-2009 flu seasons. Adults had greater intensity in the 2004-2005 flu season and lower intensity in the 2008-2009 flu seasons. Similar to results for the total population, several South Atlantic states had greater risk while Plains states had lower risk of seasonal intensity for both children and adults.

Across the three age group responses (i.e., total, children, adults), child seasonal intensity had a unique positive association with influenza B circulation and adult seasonal intensity had a unique positive association with H3 circulation among influenza A and proportion of the population in poverty. Also notable, both child and adult seasonal intensity had a negative association with estimated average household size, while the total seasonal intensity model had a positive effect.

### Drivers of epidemic duration

We also considered the mechanisms associated with epidemic duration, a measure of influenza disease burden that captures the number of weeks with heightened ILI activity. Better understanding of factors associated with longer epidemics might improve hospital preparedness in surge capacity and staffing needs and aid local public health departments in planning their influenza information or vaccination campaigns. Full results for a multi-season model of epidemic duration for the total population are reported in *Appendix 3,* but predicted value means appeared to be systematically under-estimated relative to the observed epidemic durations and the Pearson’s cross-correlation coefficient between the observed and predicted mean number of epidemic weeks was ***R*** = 0.71.

The 2004-2005 and 2007-2008 flu seasons had longer epidemics while the 2002-2003 and 2008-2009 seasons tended to have shorter epidemics. The Southeastern U.S. region (HHS region 4) had longer epidemics than other regions, while only five states with no geographic identity had significant group effects for the epidemic duration model. Epidemic duration had positive associations with the interaction between adult population and influenza H3 circulation, influenza B circulation, estimated average household size, population density, a proxy for prior immunity, and elderly vaccination coverage. There were negative associations with H3 circulation among influenza A, average flu season specific humidity, and proportion of the population in poverty. With regard to measurement factors, careseeking behavior and claims database coverage had strong positive associations with epidemic duration.

### Applications to surveillance

Considering the large volume and spatial resolution of our data, we sought to explore the robustness of our inference and model predictions under more realistic circumstances. Two sequences of models were designed to mimic different types of real-world sentinel flu surveillance systems *—fixed-location sentinels,* where the same sentinel locations reported data every year, and *moving-location sentinels,* where new sentinel locations are recruited each year. A third model sequence considered the specificity of inference and model predictions to certain *inclusion of historical data,* thus providing insight into the generalizability of our model to epidemic forecasting. We examine these applications for the total population seasonal intensity model, and these may also serve as a sensitivity analysis to missing observations. Ten replicates were performed for each model with missingness to generalize findings beyond that of random chance.

#### Sentinels in fixed locations

In this sequence of four models, 20, 40, 60, and 80% of randomly chosen county observations were removed across all years. The effect sizes of drivers were pulled towards zero as fewer sentinel counties reported ILI seasonal intensity, but the primary conclusions remained robust. We noted that the positive effect of care-seeking increased across most model replicates and insurance coverage shifted from no effect to a slightly positive effect as sentinel reporting declined (*Figure 4A*). Model predictions (county-season fitted values) remained quite robust relative to the complete model, even when 80% of counties were excluded (*Figure 4B*).

**Figure 4.**
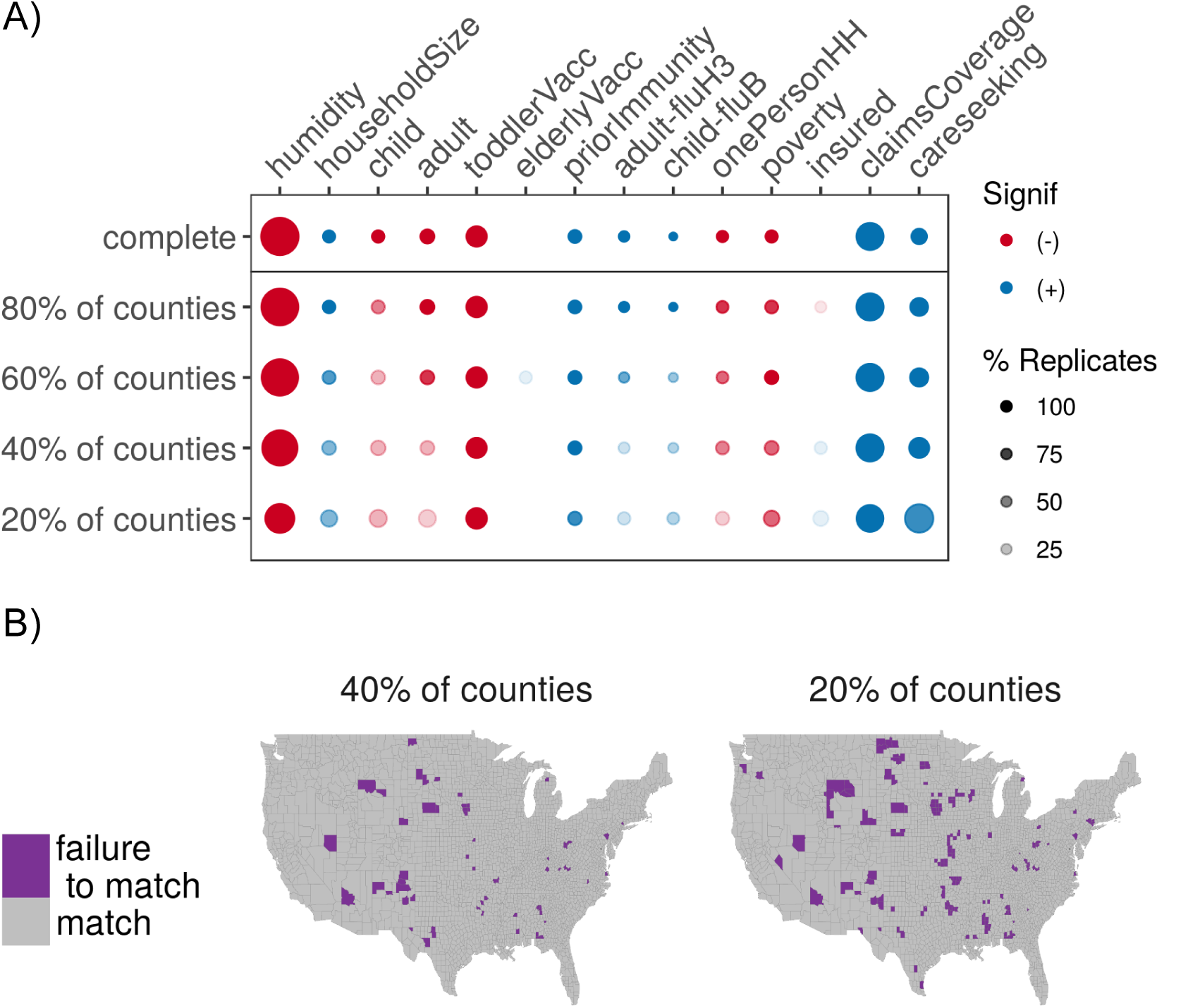
A) Diagram indicating changes to model inference as fewer fixed-location sentinels reported data. Color indicates directionality of the significant effect (blue is positive, red is negative) while greater transparency indicates a lower percentage of replicates with a significant effect (for models with missingness); dot size represents the magnitude of the posterior mean (or average of the posterior mean across replicates). Predictors with no significant effect across the sequence of models were removed for viewing ease, and absence of a dot means the effect was not significant across any replicates. B) Map of model prediction match between the complete model and the 40% and 20% reporting levels for fixed-location sentinels. Match between the complete and sentinel models were aggregated across 70 season-replicate combinations (7 seasons * 10 replicates). Color indicates match between posterior predictions in the missing and complete models (purple represents a failure to match in at least half of season-replicate combinations). **Figure 4-Figure supplement 1.** Diagram indicating changes to model inference as fewer moving-location sentinels reported data. **Figure 4-Figure supplement 2.** Map of model prediction match between the complete model and the 60% and 80% missing levels for moving-location sentinels. **Figure 4-Figure supplement 3.** Diagram indicating changes to model inference as historical seasons were randomly removed from the model. **Figure 4-Figure supplement 4.** Map of model prediction match between the complete model and models missing one, three, or five historical flu seasons.

#### Sentinels in moving locations

In this sequence of four models, 20, 40, 60, and 80% of randomly chosen seasonally-stratified observations were removed. Similar to the fixed-location sequence, drivers were pulled towards zero as fewer sentinel counties reported ILI, the drivers with the smallest means were pulled towards zero and predictors with no effect in the complete model were found to be significant (*Figure 4* supplement). Model predictions had good agreement with the complete model up to a threshold between 60 and 80% missingness, where many county-season fits suddenly became poor.

#### Inclusion of historical data

In this sequence of models, one, three, and five out of seven flu seasons in the study period were completely removed. As hinted by the inconsistency of inference across seasons in the single season model results (*Figure 3*), important drivers changed substantially when more than one season was removed, particularly when they had small effect sizes in the complete model (*Figure 4* supplement). Notably, medical claims coverage and care-seeking were two of three predictors that remained consistent in the magnitude and direction of inference across all model replicates. Model predictions were robust relative to the complete model only when one season was removed. Beyond that, many seasonal fitted values were poor, particularly for some seasons where data had been removed.

## Discussion

Using hierarchical modeling approaches, we explored the contributions of 19 potential predictors towards county-level variation in influenza disease burden across the United States during flu seasons from 2002-2003 to 2008-2009. To our knowledge, this is the first large-scale study to compare the relative importance of environmental, demographic, and socioeconomic hypotheses about influenza disease burden in addition to data reporting biases. The fine spatial resolution and high coverage of our medical claims data (estimated to represent 20% of all health care visits across the United States in our study period) enabled the comparison of multiple hypotheses, and the inclusion of several flu seasons and sensitivity analyses enhance confidence in the robustness of our findings.

Our model results suggest that South Atlantic states may experience flu seasons most acutely because they have higher seasonal intensities relative to their baselines, and greater examination of flu season surveillance and surge capacity in these areas may be warranted. We also found that a mixture of factors explained the variation in our model and that these factors changed across different cross-sections of time, thus highlighting the necessity of cross-disciplinary approaches (e.g., from sociology to epidemiology to immunology) in future pursuits of this question. Moreover, the declining importance of claims database coverage (i.e., population representativeness of the data) as coverage increased underscores the relevance of collecting and using metadata when making epidemiological inference from opportunistic sources or undesigned observational samples. The ability for our model to project relatively accurate fitted values across increasingly missing data suggests that routine sentinel surveillance in fixed locations may be more accurate for interpolating ILI disease burden among uncovered areas than surveillance across changing locations, even when fewer locations may be surveyed.

Prior studies have reported relationships between low absolute humidity and greater influenza transmission and survival in experimental settings, and that fluctuations in absolute humidity may explain the seasonality of influenza across large geographic scales (*Tamerius et al.,* 2011; *Lowen and Steel, 2014*). Our study adds to this literature in finding strong negative associations between absolute humidity and both seasonal intensity and epidemic duration. In addition, our results elucidate the debate about whether influenza transmits primarily through frequency-or density-dependent contact. Greater seasonal intensity was associated with populations with larger household sizes (a proxy for infection risk from frequent contacts), while longer epidemics were associated with larger household sizes and greater population density. We suspect that density-dependent transmission explained differences in epidemic duration but not seasonal intensity because the calculation for seasonal intensity accounted for population size; population density did not explain variation in the risk of seasonal intensity after adjusting for greater transmission among larger populations.

Household studies of influenza transmission often examine age-specific risks of household influenza introduction (*Cauchemez et al., 2004; Lau et al., 2015*), and differences in contact and travel patterns between children and adults have led to the hypothesis that children drive local transmission while adults drive global influenza spread (*Apolloni et al., 2013; Viboud et al., 2006*) Contrary to these hypotheses, larger child and adult population proportions were both associated with lower seasonal intensity. Rather than serving as proxies for local and global transmission, the complement of these predictors together may in fact capture the “high-risk” population proportion in a given location—infants, toddlers, and the elderly—which typically experience greater clinical severity (*Thompson et al., 2006*) and have higher rates of care-seeking (*Biggerstaff et al., 2012*). In examining seasonal intensity models for the child and adult populations specifically, we were surprised to find negative associations with population density and average household size, when there was no effect or a positive effect in the total population model (*Appendix 2*). While it may be that children and adults in less connected areas have greater seasonal intensity relative to their ILI baselines, these patterns may also be an artifact of smaller volumes of data among age groups.

The positive association between influenza A/H3 and adult intensity and influenza B and child intensity corroborate the results of previous epidemiological studies (*Hayward et al., 2014; Beauté et al., 2015*) and agree with the positive effect of the interaction terms between children and influenza B and adults and influenza A/H3 from our total seasonal intensity models (*Appendix 2*). Despite a positive linear correlation between the seasonal intensity and epidemic duration measures (*Appendix 4*), influenza B circulation uniquely indicated longer epidemics, in line with hypotheses that flu seasons are elongated when influenza B resurges among children after a first wave of influenza A (*Hayward et al., 2014; Beauté et al., 2015*). We acknowledge that our findings may be specific to our study period; recent research highlights the importance of childhood hemagglutinin imprinting on immune responses to subsequent influenza infections (*Gostic et al., 2016*).

We were surprised to observe that higher estimated prior immunity was associated with greater seasonal intensity and longer epidemic durations for the multi-season models and most seasons in the single-season models (some years experienced no effect). One possible interpretation is that some locations always tend to have high disease burden relative to their epidemic baselines. Prior work suggests that larger epidemics induce more antigenic drift in subsequent seasons (*Boni et al., 2004*); building off this finding, we suggest that influenza drift renews population susceptibility every flu season, even on small spatial scales. We also acknowledge limitations underlying the calculation of this predictor; in using the seasonal intensity measure to represent the previous flu season’s attack rate, we ignore asymptomatic infection, vaccination rates, and the reporting biases found to be an important component to data observation. Additionally, membership in the same antigenic cluster is a simplification of the immunity conferred by infection with a given strain. Beyond “pre-existing immunity”, we report mixed findings on the effect of flu vaccination. While higher vaccination coverage among toddlers was associated with lower seasonal intensity, we note that higher vaccination coverage among elderly was associated with longer epidemics. We posit that vaccination campaigns among elderly populations may increase in anticipation of large or severe flu seasons, due to their risk of severe complications from flu and clustered living in nursing homes.

Our study found that locations with greater poverty had lower influenza disease burden, in contrast with ample evidence that there are heightened rates of influenza-related hospitalizations, influenza-like illness, respiratory illness, neglected chronic diseases, and other measures of poor health among populations with greater material deprivation (*Hadler et al., 2016*; *Monto and Ullman, 1974*; Tam et al., 2014; *Biggerstaff et al.,* 2014b,a; *Charland et al., 2011*;*Hotez, 2008;Adler and Newman, 2002; Steptoe and Feldman, 2001*). Several possible non-exclusive explanations for this discrepancy exist. Differences in socio-economic background may change recognition and therefore reporting of disease symptoms (*Monto and Ullman, 1974*). Material deprivation and lack of social cohesion have also been implicated in lower rates of health care utilization for ILI, which would reduce the observation of influenza disease burden in our medical claims data among the poorest populations (*Charland et al., 2011; Biggerstaff et al., 2014a*). Indeed, higher rates of health care-seeking were associated with greater disease burden, while hospitals per capita had no effect among our results, which further suggests that patient-side needs and concerns captured ILI variation better than deficits in health resource availability. Future studies focused on estimation and surveillance of influenza disease burden should consider collecting and incorporating data on health care utilization in their populations of interest in order to account for reporting biases and limited forecasting ability in poorer neighborhoods (*Scarpino et al., 2016*).

Building off mechanistic explanations for measurement biases, we noted that the positive explanatory effect of claims database coverage declined as coverage itself increased throughout our study period *(Appendix 5*). Conversely, when we artificially removed counties from our model (fixed-location sentinels) or subset our data into age groups, health care-seeking behavior more strongly explained the variation in seasonal intensity among the remaining observations. These two results together suggest that statistical inference from opportunistic data samples may avoid some types of reporting biases when the coverage or volume of data achieves a minimum threshold, in response to concerns posed in *Lee et al. (2016)*. In our specific case, increases to claims database coverage or care-seeking behavior might reduce reporting biases by increasing the representativeness of a given location’s sample. Additionally, we present the concept of a network of *sentinel locations,* in contrast to sentinel physicians or hospitals, which may be composed of administrative units (e.g., counties) that were chosen for either their representativeness of the larger population or their status as an outlier (e.g., match or failure to match locations in *Figure 4,* respectively). Given the growing availability of health-associated big data in infectious disease surveillance (*Bansal et al., 2016; Simonsen et al., 2016*), we project the possibility that sentinel locations may report high volume digital health data from disparate sources to a central public health organization and that the informed choice of sentinels may improve the robustness of sentinel surveillance systems.

We urge caution in the interpretation of our results because they are correlative and prone to invoking the ecological fallacy, where statistical inference about a group (in our case, county populations) is falsely assumed to apply at the individual level (*Morgenstern, 1982; Robinson, 2009*). Future research should build off our study to design experiments that may provide causal or individual-level evidence that supports or rejects these hypotheses. We also acknowledge the limitations of the spatial and temporal resolutions of the data used in our analysis. Previous work suggests that statistically-identified drivers of disease distributions depend on the spatial scale of analysis (*Cohen et al., 2016*), and our results may be biased by the county unit observations of our disease data. In addition, we incorporated multiple scales of predictors (county, state, and HHS region) according to the best available data, thus potentially altering our statistical inference, although we did attempt to account for differences in variation across these different predictors with the inclusion of group effects. In addition, we note that the nature of our disease burden estimation procedure means that a given county’s seasonal intensity is relative to its own baseline across years. It may not be appropriate to use our model predictions to inform national-level decision makers about absolute intensity of the flu season in a given location, although local public health departments could use our procedure to assess intensity in a given year relative to that of previous flu seasons.

## Methods

### Medical claims data

Weekly visits for influenza-like illness (ILI) and any diagnosis from October 2002 to May 2009 were obtained from a records-level database of CMS-1500 US medical claims managed by IMS Health and aggregated to three-digit patient US zipcode prefixes (zip3s), where ILI was defined with International Classification of Diseases, Ninth Revision (ICD-9) codes for: direct mention of influenza, fever combined with respiratory symptoms or febrile viral illness, or prescription of oseltamivir. Medical claims have been demonstrated to capture respiratory infections accurately and in near real-time (*Cadieux and Tamblyn, 2008*; *Santillana et al., 2016*), and our specific dataset was validated to independent ILI surveillance data at multiple spatial scales and age groups and captures spatial dynamics of influenza spread in seasonal and pandemic scenarios (*Viboud et al., 2014; Gog et al., 2014; Charu et al., 2017*).

We also obtained database metadata from IMS Health on the percentage of reporting physicians and the estimated effective physician coverage by visit volume; these data were used to generate “measurement” predictors (*Table* 1). ILI reports and measurement factors at the zip3-level were redistributed to the county-level according to population weights derived from the 2010 US Census ZIP Code Tabulation Area (ZCTA) to county relationship file, assuming that ZCTAs that shared the first three digits belonged to the same zip3.

### Defining influenza disease burden

We performed the following data processing steps for each county-level time series of ILI per population: i) Fit a LOESS curve to non-flu period weeks (flu period defined as November through March each year) to capture moderate-scale time trends (span = 0.4, degree = 2); ii) Subtract LOESS predictions from original data to detrend the entire time series; iii) Fit a linear regression model with annual harmonic terms and a time trend to non-flu period weeks (*Yu et al., 2013*); iv) Counties “had epidemics” in a given flu season if at least two consecutive weeks of detrended ILI observations exceeded the ILI epidemic threshold during the flu period (i.e., epidemic period) (*Denoeud et al., 2007*). The epidemic threshold was the upper bound of the 95% confidence interval for the linear model prediction. Counties with a greater number of consecutive weeks above the epidemic threshold during the non-flu period than during the flu period were removed from the analysis.; v) Disease burden metrics were calculated for counties with epidemics.

Multiple measures of influenza disease burden were defined for each county. For a given season: *seasonal intensity* was the one plus the sum of detrended ILI observations during the epidemic period (shifted by one to accomodate the likelihood distribution); *epidemic duration* was the number of weeks in the epidemic period and counties without epidemics were assigned the value zero.

### Predictor data collection and variable selection

Quantifiable proxies were identified for each hypothesis found in the literature, and these mechanistic predictors were collected from probability-sampled or gridded, publicly available sources and collected or aggregated to the smallest available spatial unit among US counties, states, and Department of Health and Human Services (HHS) regions for each year or flu season in the study period, as appropriate (*Table 1*, *Appendix 5*).

We selected one predictor to represent each hypothesis according to the following criteria, in order: i) Select for the finest spatial resolution; ii) Select for the greatest temporal coverage for years in the study period; iii) Select for limited multicollinearity with predictors representing the other hypotheses, as indicated by the magnitude of Spearman rank cross-correlation coefficients between predictor pairs. We also compared the results of single predictor models and our final multivariate models as another check of multicollinearity (*Appendix 5*). For the modeling analysis, if a predictor had missing data at all locations for an entire year, data from the subsequent or closest other survey year were replicated to fill in that year. If a predictor data source was available only at the state or region-level, all inclusive counties were assigned the corresponding state or region-level predictor value (e.g., assign estimated percentage of flu vaccination coverage for state of California to all counties in California). Predictors were centered and standardized prior to all exploratory analyses and modeling, as appropriate. Interaction terms comprised the product of their component centered and standardized predictors. Data cleaning and exploratory data analysis were conducted primarily in R (*R Core Team, 2015*). Final model predictors are described below, and our hypotheses for each predictor are described in *Table 1.*

#### Environmental data

Daily specific humidity data on a 2m grid were collected from the National Oceanic and Atmospheric Administration (NOAA) North American Regional Reanalysis (NARR), provided by the NOAA/OAR/ESRL PSD, Boulder, Colorado, USA, from their website at http://www.esrl.noaa.gov/psd/. Values were assigned to the grid point nearest to the county centroid.

Readings of fine particulate matter, defined as pollutants with aerodynamic diameter less than 2.5 micrometers, were collected from the CDC WONDER database at the county and daily scales from their website at https://wonder.cdc.gov/.

#### Social contact and population data

Annual total and age-specific population data were taken from the intercensal population estimates and land area and number of housing units were reported during the 2000 and 2010 Census; both datasets were available at the county scale from the U.S. Census Bureau. These data were used to calculate proportion of total population that are children (5-19 years old) and adults (20-69 years old), population density by land area, and estimated average household size.

#### Flu-specific data

Annual flu vaccination rates for toddlers (19-35 months old) and the elderly (≥ 65 years old) were estimated at the state-level from the Centers for Disease Control and Prevention (CDC) National Immunization Survey and Behavioral Risk Factor Surveillance System, respectively. Annual proportion of A-typed flu samples subtyped as H3 and annual proportion of confirmed flu samples typed as B across U.S. Department of Health and Human Services (HHS) regions were collected by WHO/NREVSS Collaborating Labs and available at the CDC FluView website at http://www.cdc.gov/flu/weekly/fluviewinteractive.htm.

#### Prior immunity

For a given county, a proxy for prior immunity was derived from the following data: 1) the previous flu season’s total population seasonal intensity; the proportion of positive flu strains identified as A/H3, A/H1, and B in the broader HHS region during 2) the previous flu season and 3) the current flu season; 4) the most prominently circulating flu strain for each category (A/H3, A/H1, or B) for each flu season; 5) antigenic clusters for A/H3 and A/H1 strains as identified in *Du et al. (2012)*; *Liu et al.* (*2015*); and 6) Victoria-or Yamagata-like lineages for B strains as noted in *Bedford et al.* (*2014*). Data for items 1-3 are described above in “Defining influenza disease burden” and “Flu-specific data.” We obtained the antigenic characterizations for circulating strains (item 4) from CDC influenza season summaries, which are available at https://www.cdc.gov/flu/weekly/pastreports.htm.

Using these data, we calculated a proxy of prior immunity that captures “the proportion of individuals infected in the previous flu season that would have protection during the current flu season, accounting for the distribution of circulating flu strains.” For each flu category among A/H3, A/H1, and B, we calculated the product of the previous and current year’s proportion of total circulation and a binary value to indicate if previous and current strains were from the same antigenic cluster or lineage (1 = same cluster/lineage, 0 = different cluster/lineage). For a given county, these products were summed across A/H3, A/H1, and B, and multiplied by the previous year’s seasonal intensity.

#### Socioeconomic and access to care data

Annual data on number of hospitals were obtained at the county-level from the Health Resources and Services Administration (HRSA) Area Health Resources Files (AHRF). County-level data on proportion of households with a single person were obtained from five-year averages of American Community Survey (ACS) estimates, which were available starting in 2005. Annual estimates on proportion of the population in poverty was obtained at the county-level from the model-based Small Area Income and Poverty Estimates (SAIPE). Annual estimates on proportion of the population with health insurance was obtained at the county-level from the model-based Small Area Health Insurance Estimates (SAHIE). SAIPE and SAHIE are both products of the U.S. Census Bureau that were derived from the Current Population Survey or ACS.

#### Medical claims measurement factors

IMS Health provided us with weekly aggregated data on visits for any diagnosis by age group and location. Care-seeking behavior was defined as the total visits per population size from November through April of a given flu season. Claims database coverage was the estimated physician coverage among all physicians registered by the American Medical Association in the IMS Health medical claims database.

### Model structure

We present the most common version of our model structure here. The generic model for county-year observations (for *i* counties and *t* years) of influenza disease burden *y_it_* is:

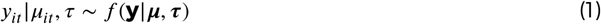

where **y** = (*y*_1_,…, *y_n_*)*’* denotes the vector of all observations (*Equation 1*). We modeled the mean of the observed disease burden magnitude (*μ_i_*), where *f*(**y***|μ, τ*) is the distribution of the likelihood of the disease burden data, parameterized with mean *μ =* (*μ_1_,…, μ_n_*)′ and precision τ, as appropriate to the likelihood distribution (N.B., for the Poisson likelihood, *μ = 1/τ*)

The mechanisms driving disease burden were modeled:

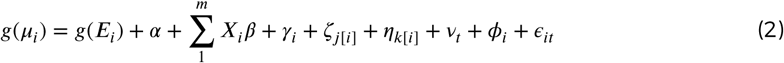

where *g*(.) is the link function, *α* is the intercept, there are *m* socio-environmental and measurement predictors (*X_i_*’s), and *E_i_* is an offset of the expected disease burden, such that *Equation 2* models the relative risk of disease (*μ_i_/Ε_i_*) in county *i,* common in disease mapping (*Lawson, 2013;Banerjee et al., 2015; Waller and Carlin, 2010).* Group terms at the county, state, region, and season levels (*γ_i_,ζ_j_*_[*i*]_*,η_k[i]_,ν_t_,* respectively) and the error term (ϵ_*it*_) are independent and identically distributed (*iid*)

Geographical proximity appears to increase the synchrony of flu epidemic timing (*Schanzer et al., 2011b; Stark et al., 2012*), while connectivity between cities has been linked with spatial spread in the context of commuting and longer distance travel (*Charaudeau et al., 2014;Brownstein et al., 2006; Crépey and Barthélemy, 2007; Lemey et al., 2014*) We modeled county spatial dependence *φ_i_* with an intrinsic conditional autoregressive (ICAR) model, which smooths model predictions by borrowing information from neighbors (*Besag et al., 1991*):

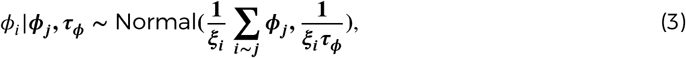

where *ξ_i_* represents the number of neighbors for node *i. ϕ_j_* is a vector indicating the neighborhood relationship between node *i* and all nodes *j* (*i* ~ *j*) and *τ_ϕ_* is the precision parameter (*Equation 3*).

### Model fit, sensitivity, and validation

To assess model fit, we examined scatterplots and Pearson’s cross-correlation coefficients between observed and fitted values for the relative risk of total population seasonal intensity and for epidemic duration. We also examined scatterplots of standardized residuals and fitted values; standardized residuals were defined as (*y – μ_ŷ_)Ισ _ŷ_,* where *μ _ŷ_* is the fitted value posterior mean and *σ _ŷ_* is the fitted value standard deviation. Model sensitivity was assessed by comparing model fits and inference robustness when observations were randomly removed from the model, as described below under “Applications to missing data & inference robustness.”

For each disease burden measure, we compared models with no spatial dependence, county-level dependence only, state-level dependence only, and both county and state-level dependence. The goal of the county-level dependence was to capture local population flows, while state-level dependence attempted to capture state-level flight passenger flows (details in *Appendix 1*). We determined that models with only county-level spatial neighborhood structure best fit the data after examining the Deviance Information Criteria (DIC) values and spatial dependence coefficients of the four model structures. County-level spatial structure was subsequently used in all final model combinations. We report results from models with county-level dependence only.

For model validation, we compared model fitted values for seasonal intensity with CDC ILI and laboratory surveillance data (details in *Appendix 1*).

### Statistical analysis

The goals of our modeling approach were to i) estimate the contribution of each predictor to influenza disease burden, ii) predict disease burden in locations with missing data, and iii) improve mapping of influenza disease burden. We performed approximate Bayesian inference using Integrated Nested Laplace Approximations (INLA) with the R-INLA package (www.r-inla.org) (*Rue et al., 2009; Martins et al., 2013*). INLA has demonstrated computational efficiency for latent Gaussian models and produced similar estimates for fixed parameters as established implementations of Markov Chain Monte Carlo (MCMC) methods for Bayesian inference (*Carroll et al., 2015*). Extensions to INLA have enabled its application to spatial, spatio-temporal, and zero-inflated models (*Lindgren et al., 2011; Arab, 2015*), which is implicated in INLA’s growing use in the disease mapping and spatial ecology communities (*Schrödle and Held, 2011; Blangiardo et al., 2013*)

Seasonal intensity was modeled with a lognormal distribution, and epidemic duration was modeled with a Poisson distribution and log link and excluded the offset term in *Equation 2.* Consequently, we note that all seasonal intensity models examine the relative risk of seasonal intensity, while epidemic duration models directly examine the duration in weeks. Multi-season models included all terms in *Equation 2,* while single-season models included all terms in *Equation 2* except the season grouping (*v_t_*). Model coefficients were interpreted as statistically significant if the 95% credible interval for a parameter’s posterior distribution failed to include zero.

### Applications to missing data & inference robustness

We considered the robustness of our total population model results by refitting models where 20%, 40%, 60% and 80% of all county observations were replaced with NAs (*sentinels in fixed locations*), and where 20%, 40%, 60% and 80% of model observations were stratified by season and randomly replaced with NAs (*sentinels in moving locations*). We also refit three models where one, three, and five of seven flu seasons were randomly chosen and completely replaced with NAs *(inclusion of historical data).* To account for variability due to random chance, models were replicated ten times each with different random seeds. For each sequence of missingness, we compared the magnitude and significance of socio-environmental and measurement drivers, and the posterior distributions of county-season fitted values. Fitted value distributions were noted as significantly different if the interquartile ranges for two fitted values failed to overlap with each other.

## Acknowledgments

The authors thank IMS Health for kindly sharing the medical claims data.

## Funding

This work was supported by the Jayne Koskinas Ted Giovanis Foundation for Health and Policy (JKTG) [dissertation support grant to ECL]; and the RAPIDD Program of the Science & Technology Directorate, Department of Homeland Security and the Fogarty International Center, National Institutes of Health. The content is solely the responsibility of the authors and does not necessarily reflect the official views of JKTG, the National Institutes of Health, or IMS Health.

## Appendix 1

### Seasonal intensity model fit and validation Model fit

**Appendix 1 Figure 1.**
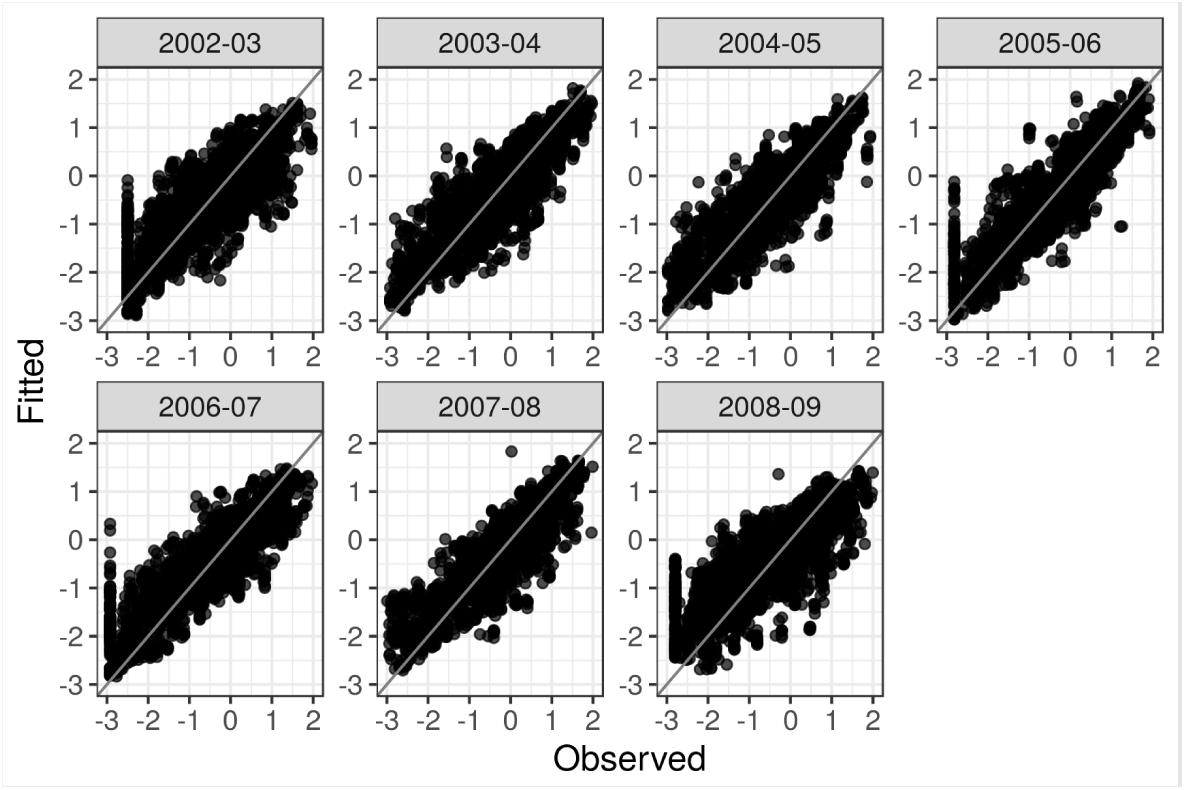
Observed vs. fitted values for the relative risk of total population seasonal intensity.

**Appendix 1 Figure 2.**
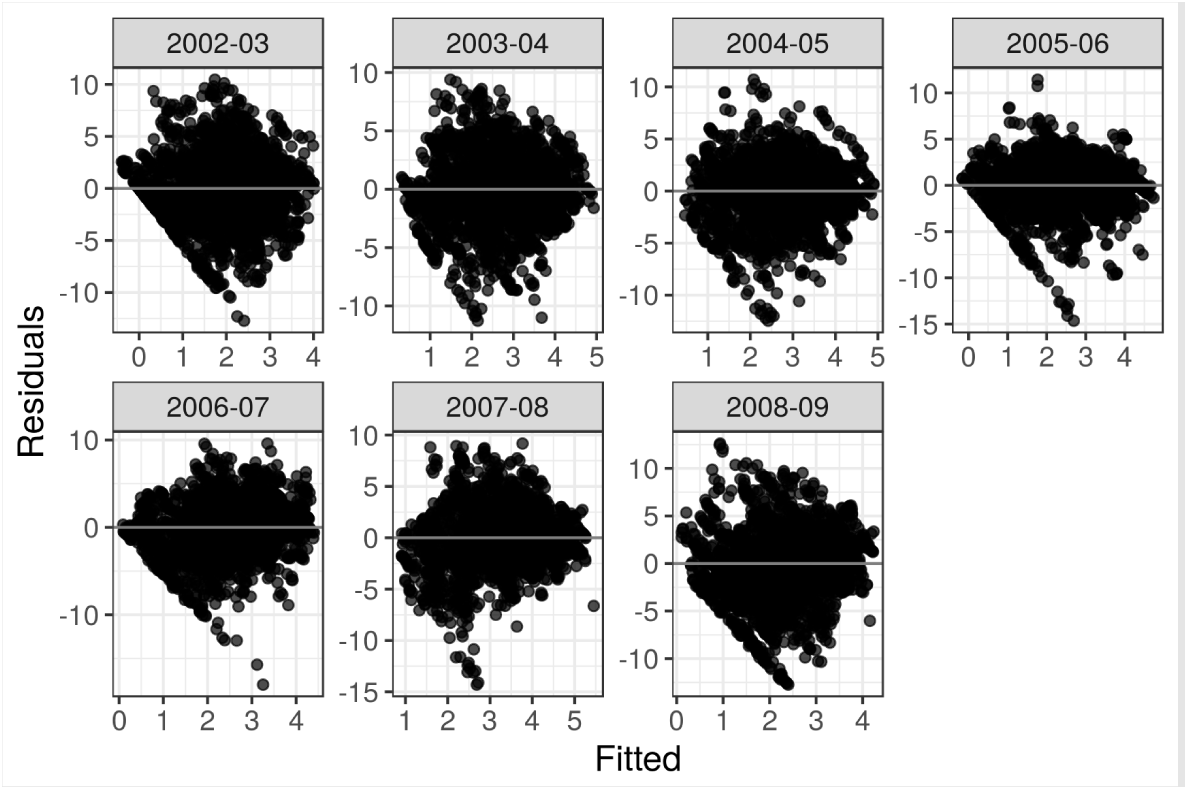
Residuals vs. fitted values for the total population log seasonal intensity.

### Selection for spatial dependence terms

To determine county-level spatial neighbors, we started with the 2010 U.S. Census Bureau 500k resolution county shapefile, and connected abutting counties that were separated by bodies of water. We then used the clean shapefile to identify neighbors as counties that shared borders.

To define state-level spatial neighbors, monthly air travel passenger flows were collected from the Bureau of Transportation Statistics T-100 Domestic Market (U.S. Carriers) table from their website at http://www.transtats.bts.gov/. Airport flows were aggregated to the state-level and states were neighbors if passengers traveled between them from November 2007 through April 2008.

**Appendix 1 Table 1.** Comparison of total seasonal intensity models with different spatial dependence structures according to Deviance Information Criterion (DIC).

| Spatial dependence structure | DIC |
| --- | --- |
| None (no $\phi$ terms) | 40,932 |
| County only (bordering neighbors, $\phi_i$) | 40,070 |
| State only (flight passenger flows, $\phi_j$) | 40,933 |
| County and state together ($\phi_i$ and $\phi_j$) | 40,070 |

**Appendix 1 Figure 3.**
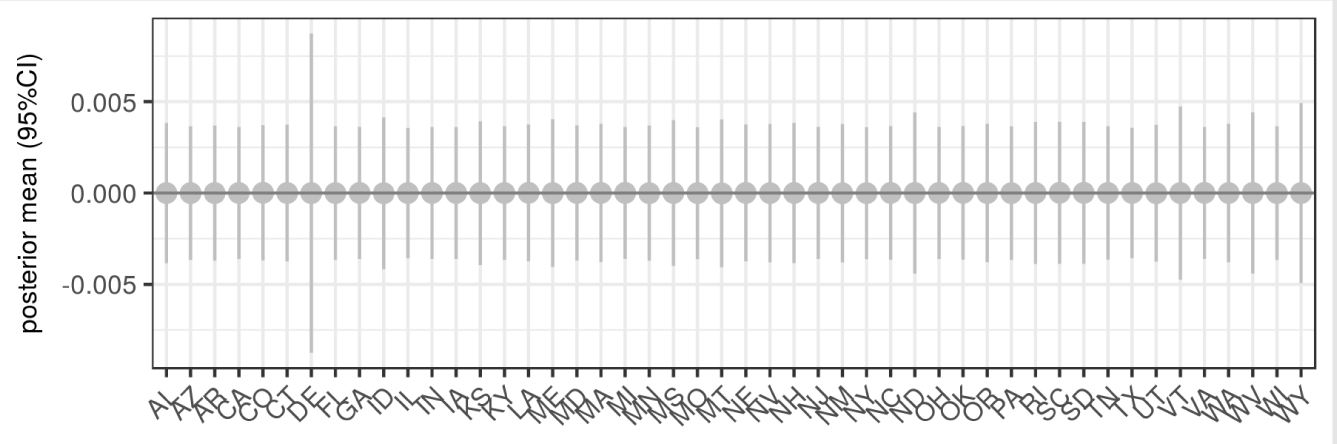
95% credible intervals for the state-level spatially structured coefficients when modeling seasonal intensity with state-level spatial dependence (*ϕ_j_*). None of the spatially structured state coefficient distribution were significant.

### Validation to CDC surveillance data

We collected a) the percentage of ILI out of all patient visits among the total population, and child and adult populations as reported by CDC’s ILINet, and b) the percentage of positive influenza laboratory confirmations as reported by CDC laboratory surveillance. = We note that child and adult ILI percentage was calculated with a denominator of patient visits across all age groups due to limited data availability. Both CDC surveillance systems were reported at the HHS region level and aggregated cumulatively for each flu season in our study period. We then examined scatterplots and Pearson cross-correlation coefficients (double-sided test where *H_0_* = no difference) between the mean model fits (where we took the mean across all counties in a given HHS region) and each CDC surveillance dataset.

**Appendix 1 Figure 4.**
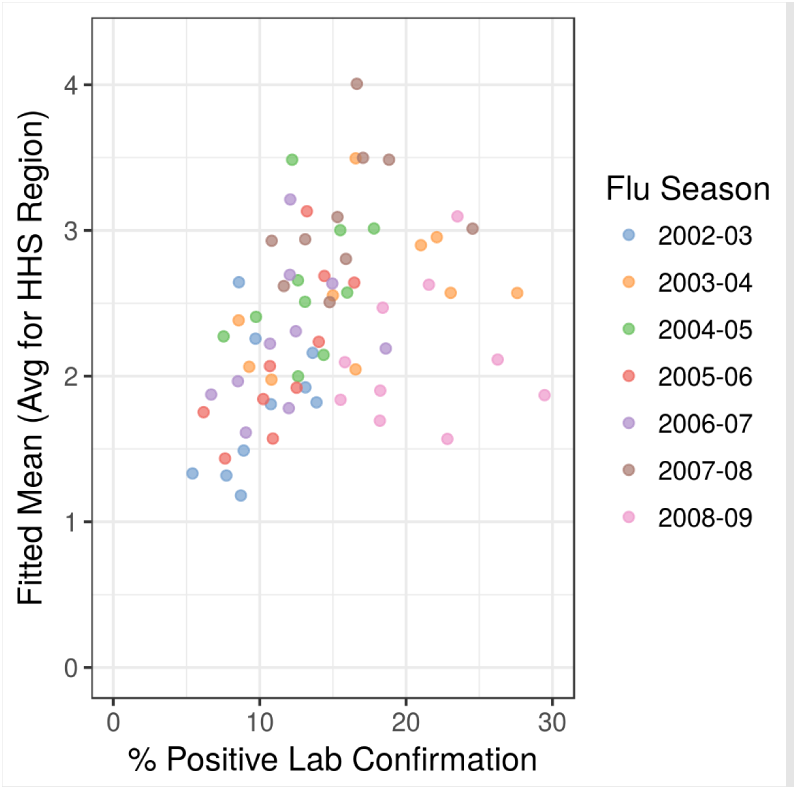
Mean model fit averaged across counties in a given HHS region vs. percentage of positive influenza laboratory confirmations in a given HHS region and flu season. The Pearson cross-correlation coefficient was 0.35 with a p-value of 0.003 for a double-sided hypothesis test.

**Appendix 1 Figure 5.**
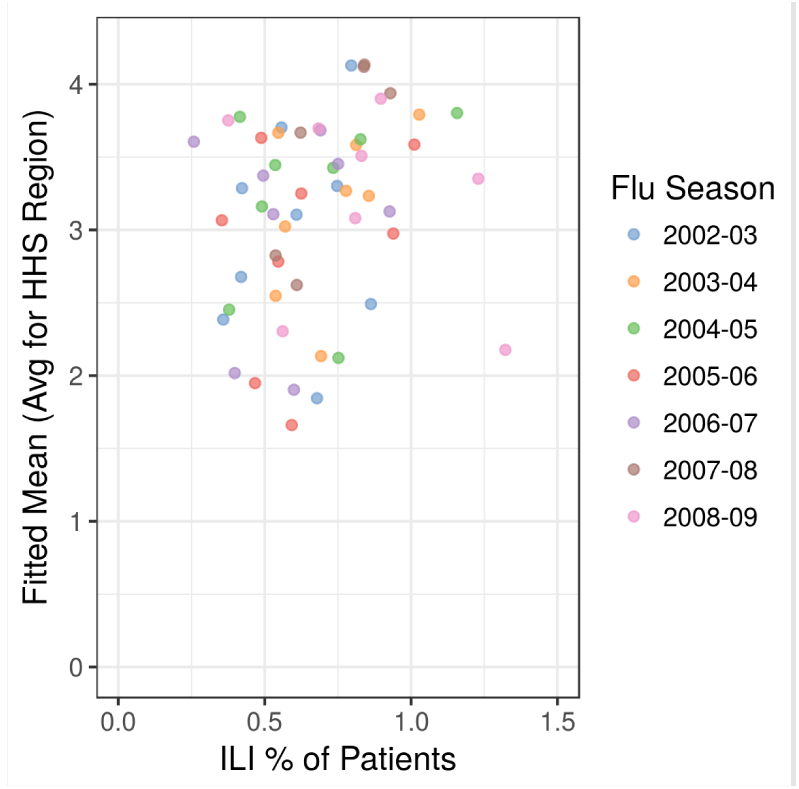
Mean model fit averaged across counties in a given HHS region vs. cumulative percentage of ILI visits in a given HHS region for all age groups. The Pearson cross-correlation coefficient was 0.38 with a p-value of 0.001 for a double-sided hypothesis test.

**Appendix 1 Figure 6.**
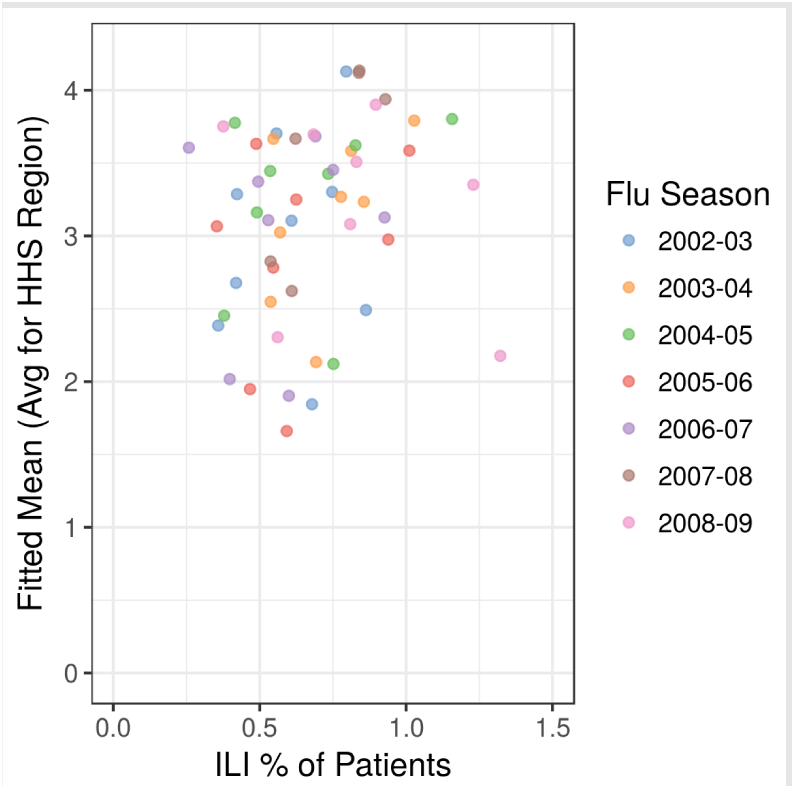
Mean model fit averaged across counties in a given HHS region vs. cumulative percentage of ILI visits in a given HHS region for children. The Pearson cross-correlation coefficient was 0.42 with a p-value of 0.0002 for a double-sided hypothesis test.

**Appendix 1 Figure 7.**
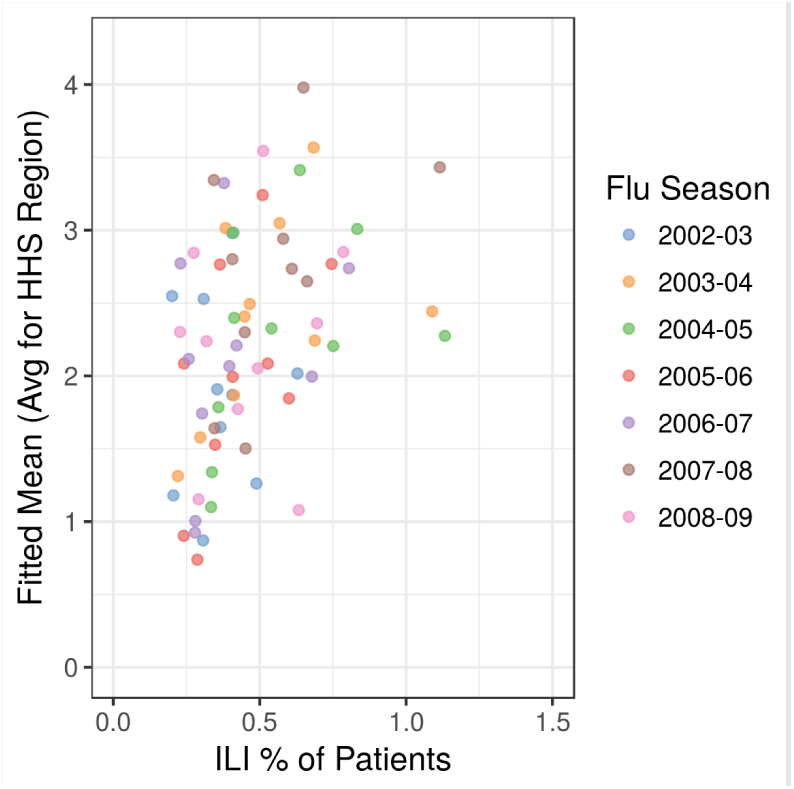
Mean model fit averaged across counties in a given HHS region vs. cumulative percentage of ILI visits in a given HHS region for adults. The Pearson cross-correlation coefficient was 0.42 with a p-value of 0.0003 for a double-sided hypothesis test.

## Appendix 2

### Age-specific drivers of seasonal intensity Model Fit

**Appendix 2 Figure 1.**
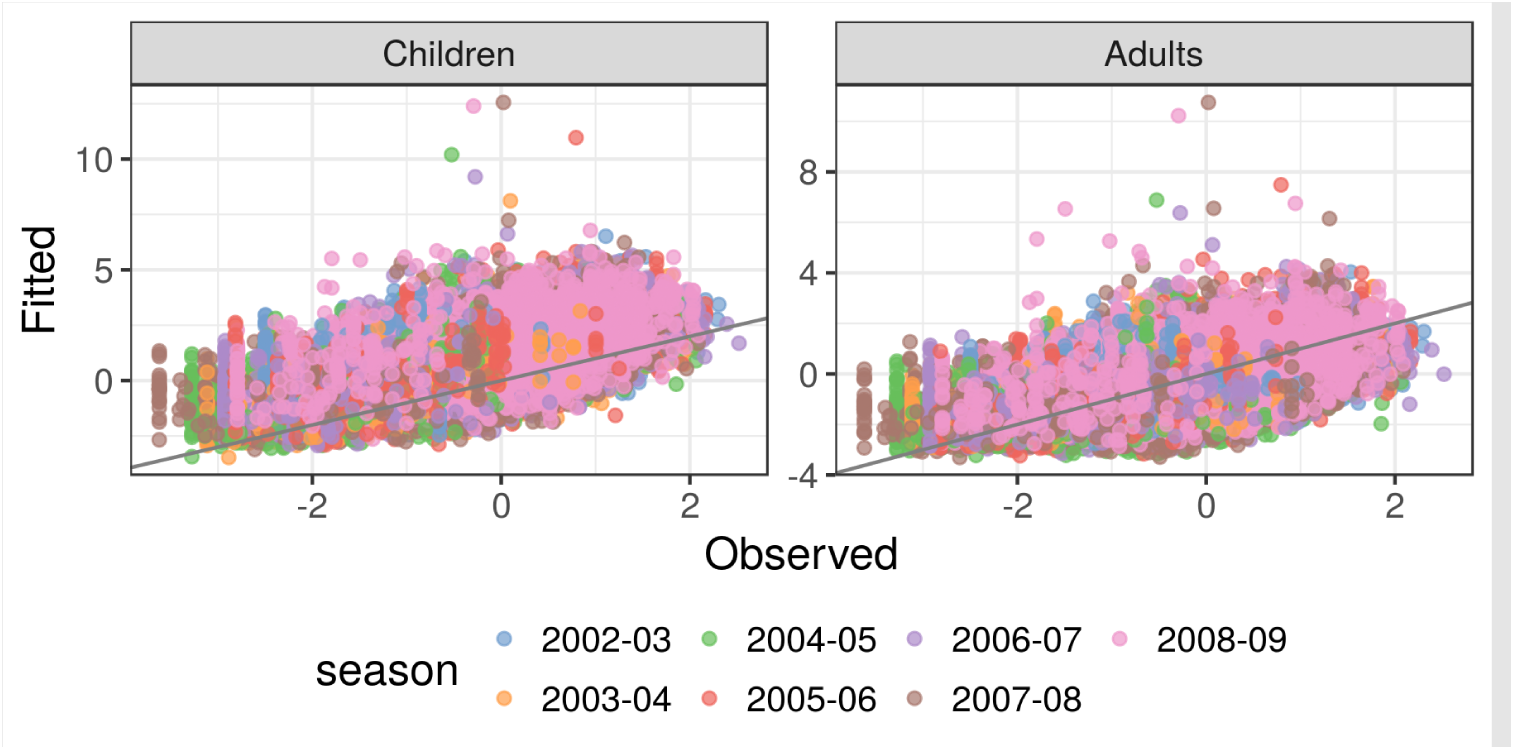
Comparison of observed and predicted relative risk of seasonal intensity across flu seasons from 2002-2003 through 2008-2009 for children and adults.

### Spatial and temporal patterns

**Appendix 2 Figure 2.**
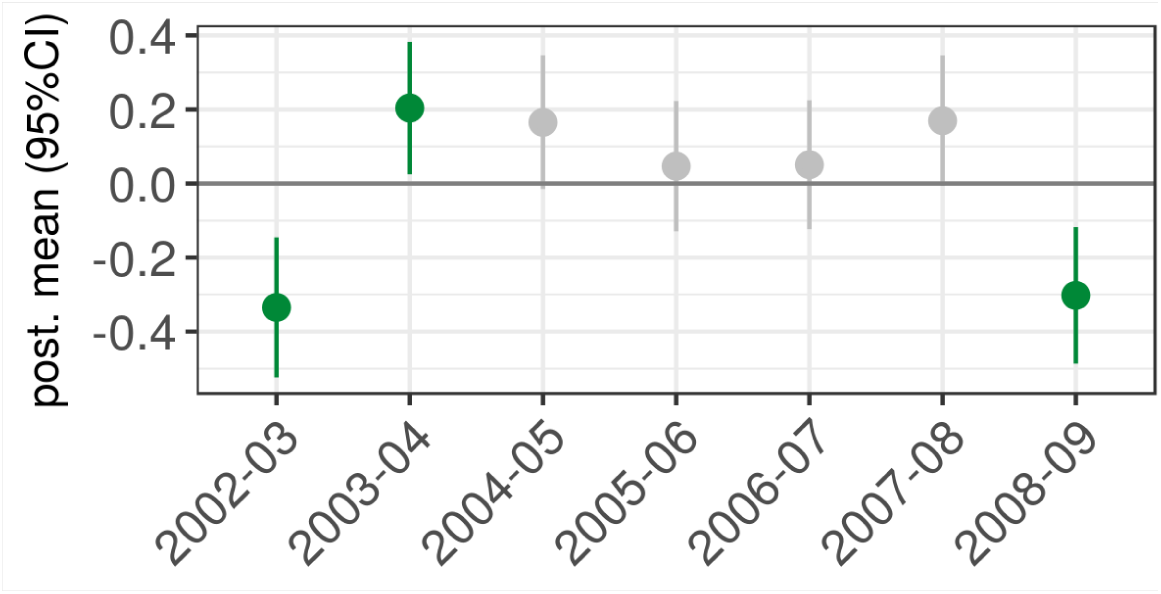
Temporal group effects for seasonal intensity among children. 95% credible interval for flu season coefficients in child population seasonal intensity.

**Appendix 2 Figure 3.**
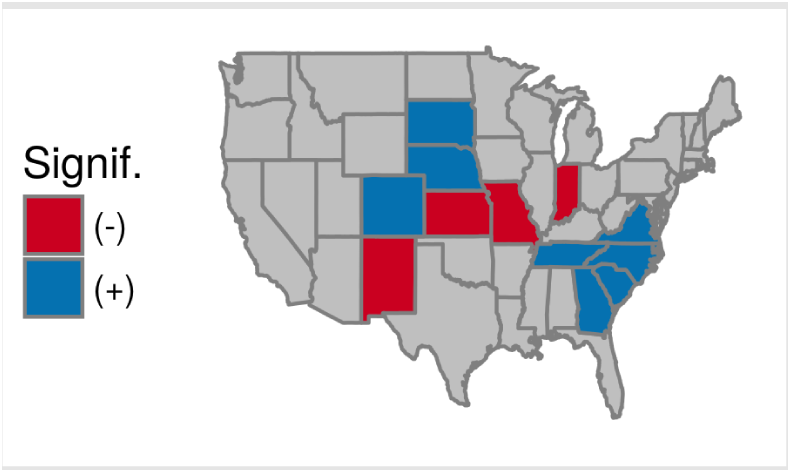
Spatial group effects for seasonal intensity among children. Continental U.S. maps highlighting states with significantly greater or lower child seasonal intensity.

**Appendix 2 Figure 4.**
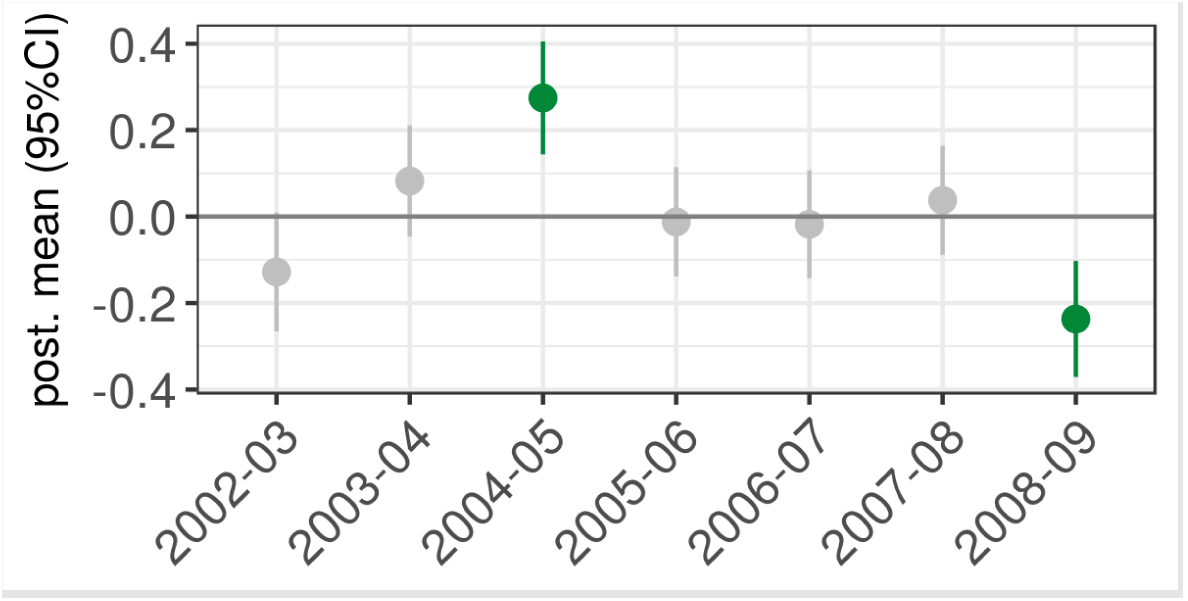
Temporal group effects for seasonal intensity among adults. 95% credible interval for flu season coefficients in adult population seasonal intensity.

**Appendix 2 Figure 5.**
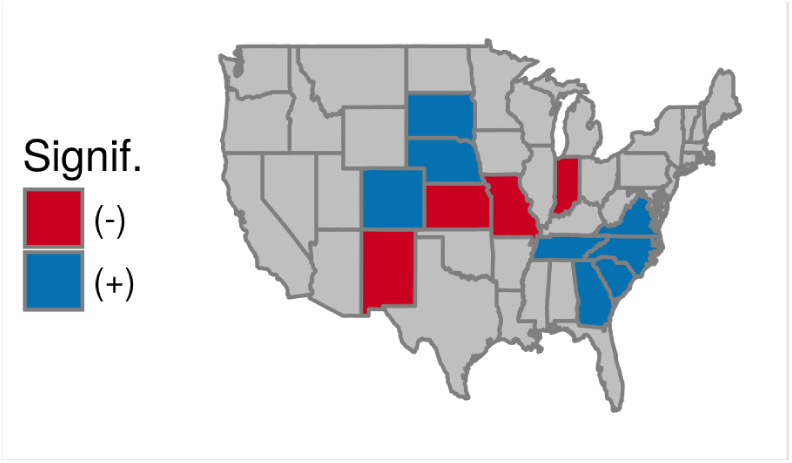
Spatial group effects for seasonal intensity among adults. Continental U.S. maps highlighting states with significantly greater or lower adult seasonal intensity.

### Socio-environmental and measurement drivers

In reference to the total seasonal intensity results, the child and adult models shared the same significant positive associations for the interaction term between child population and influenza B circulation and a proxy for prior immunity, and the same significant negative associations for adult and child population sizes, average flu season specific humidity, proportion of single person households, and infant vaccination coverage. The child and adult models shared a positive association with hospitals per capita where the total population model had no effect, and a negative association with estimated average household size where the total population model had a positive effect.

Child population seasonal intensity had a unique positive association with influenza B circulation and a unique negative association with elderly vaccination coverage. Adult population seasonal intensity had unique positive associations with H3 circulation among influenza A, proportion of the population in poverty, and elderly vaccination coverage, and a unique negative association with the interaction between adult and influenza H3.

Similar to the total population models, the child and adult seasonal intensity models had significant positive associations with careseeking behavior and claims database coverage. However, both the child and adult seasonal intensity models had significant negative associations with proportion of the population with health insurance, where the total population model demonstrated no effect.

**Appendix 2 Figure 6.**
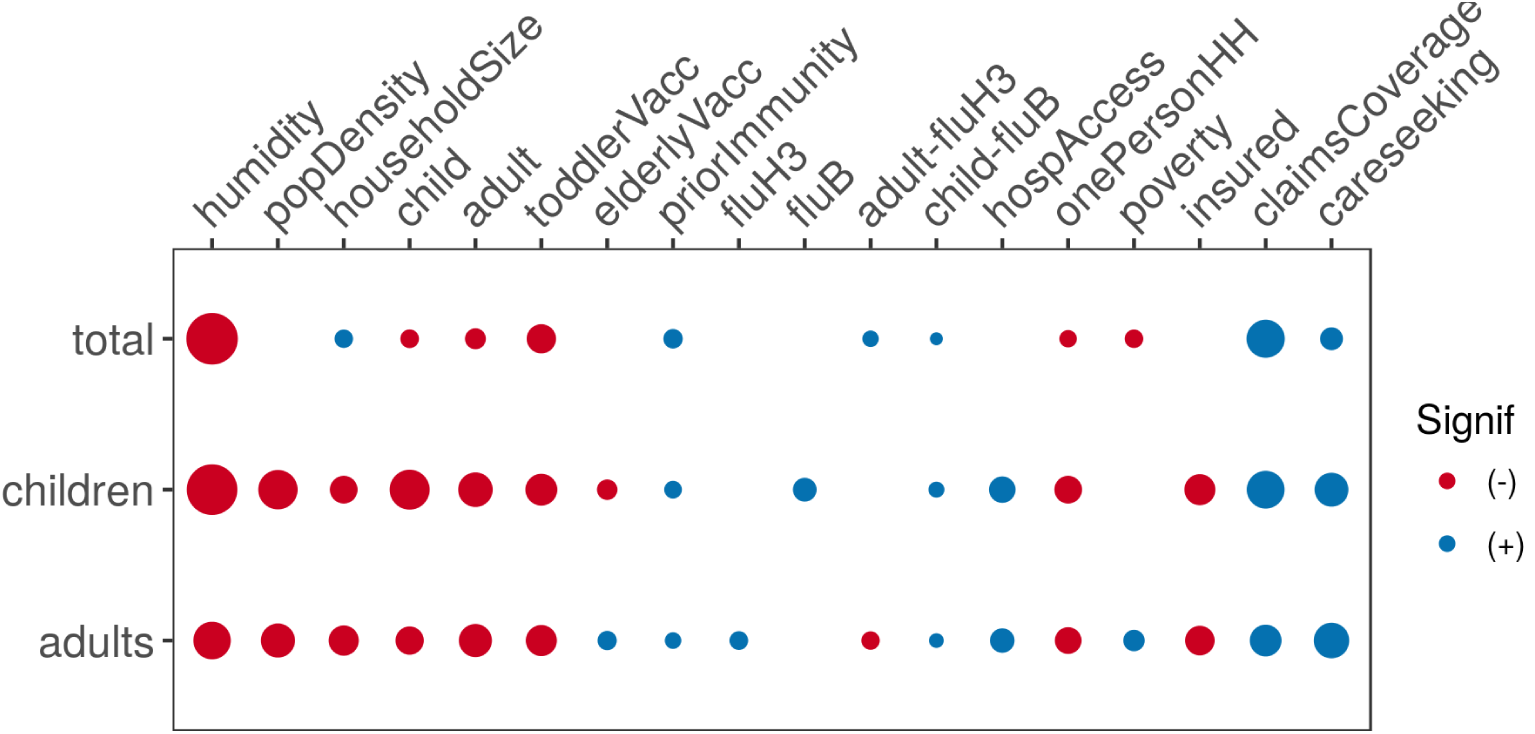
Diagram comparing model inference between total, child, and adult seasonal intensity.

## Appendix 3

### Drivers of epidemic duration Model fit

**Appendix 3 Figure 1.**
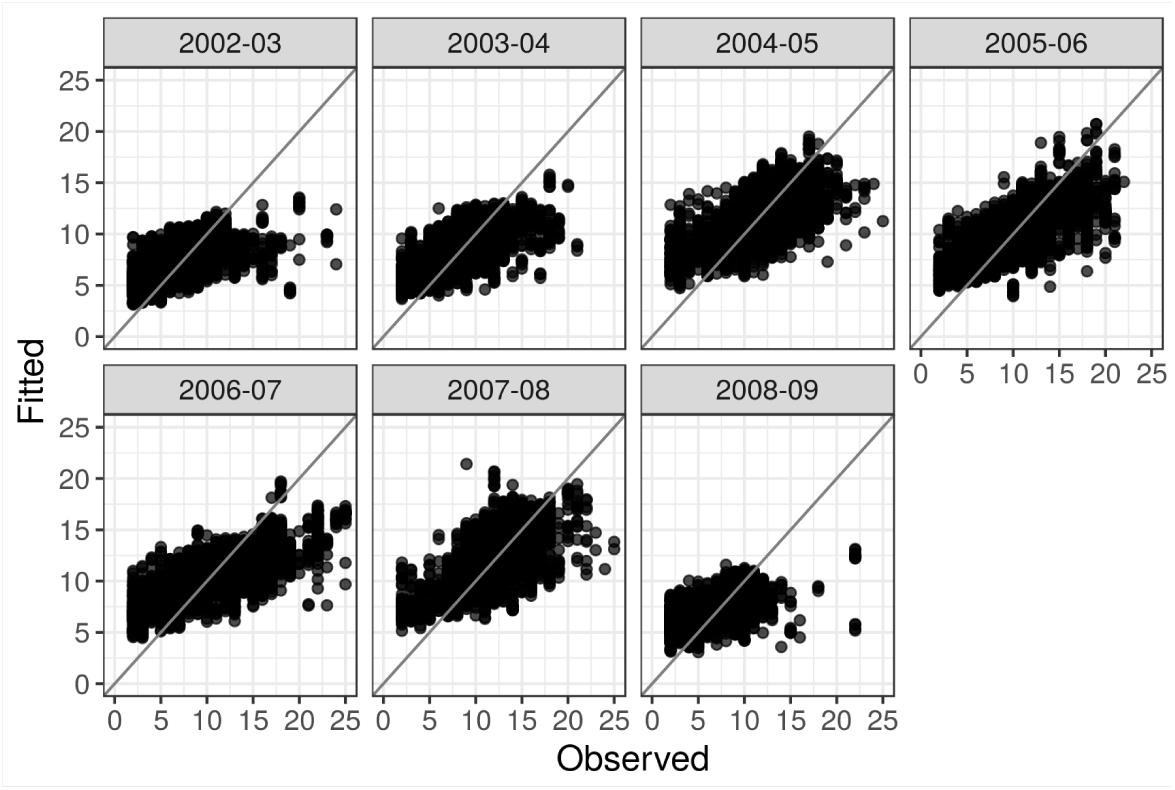
Observed versus fitted values for epidemic duration.

**Appendix 3 Figure 2.**
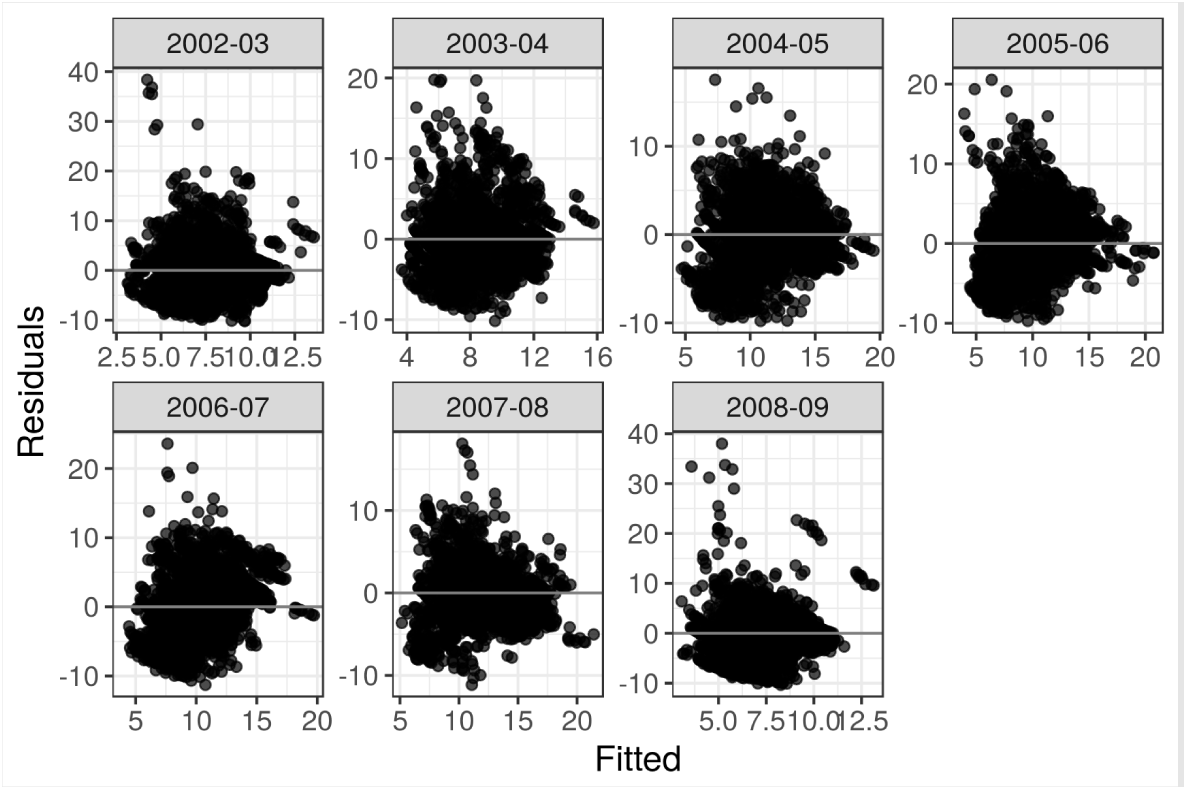
Residuals versus fitted values for epidemic duration.

**Appendix 3 Figure 3.**
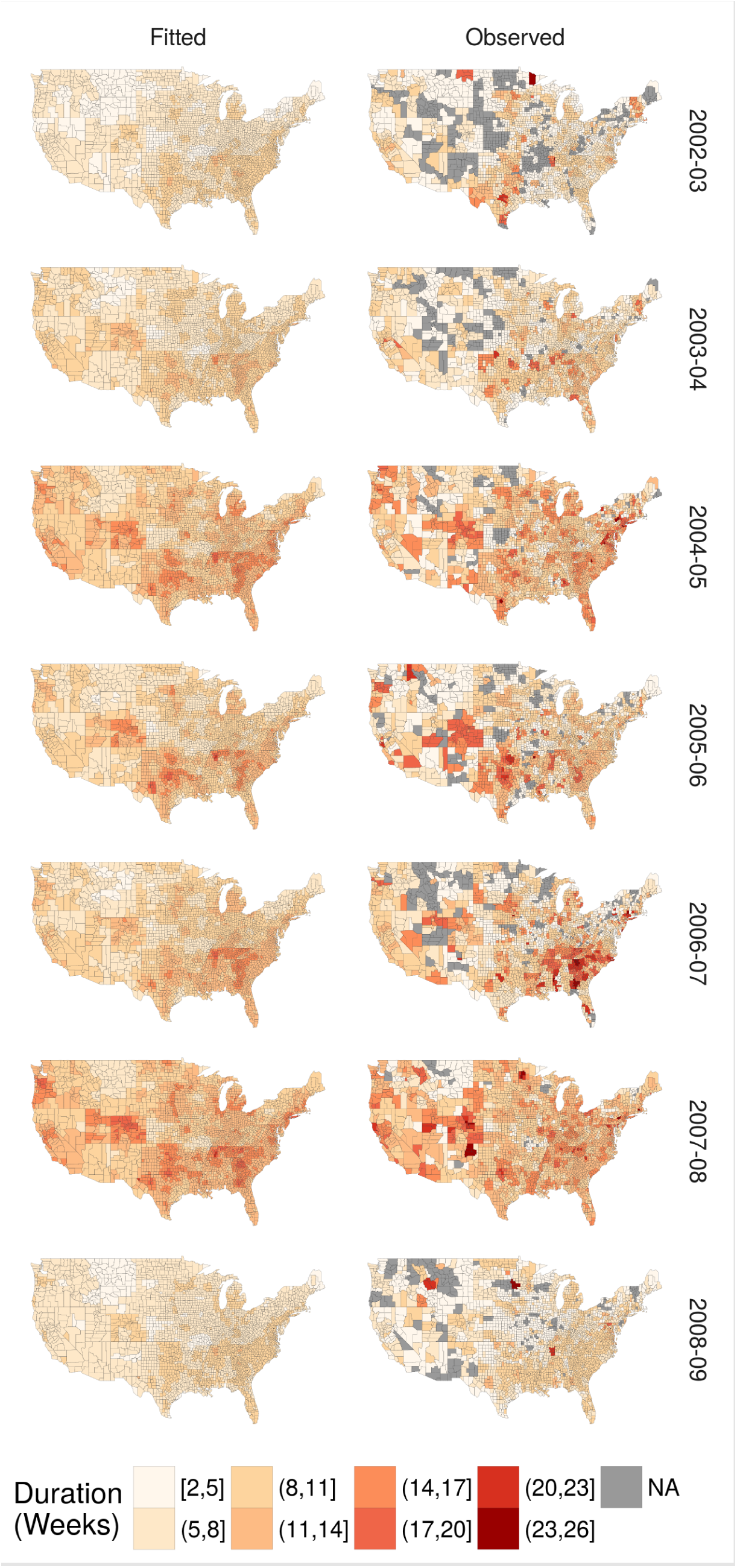
Continental U.S. county maps for fitted (left) and observed (right) epidemic duration in weeks from 2002-03 through 2008-09.

### Spatial and temporal patterns

**Appendix 3 Figure 4.**
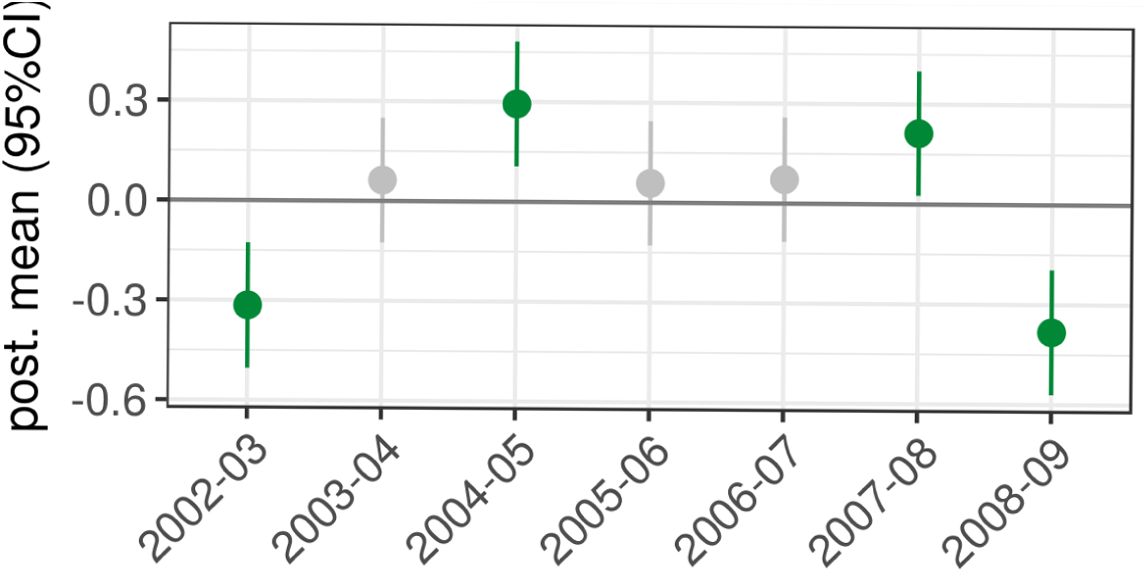
Temporal group effects for influenza-like illness. 95% credible interval for flu season coefficients in epidemic duration.

**Appendix 3 Figure 5.**
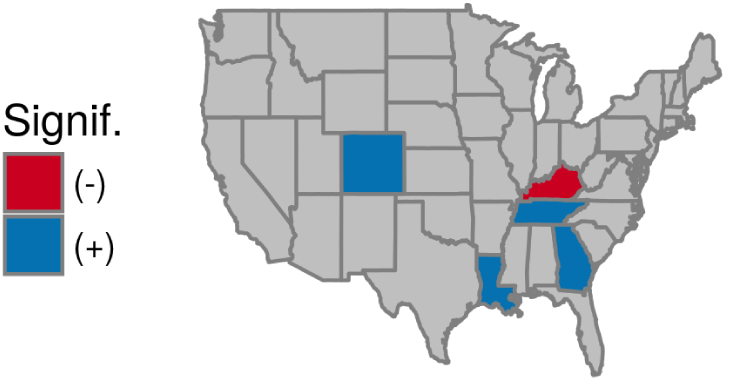
Spatial group effects for influenza-like illness. Continental U.S. map highlighting states with significantly longer or shorter epidemic durations.

### Socio-environmental and measurement drivers

**Appendix 3 Figure 6.**
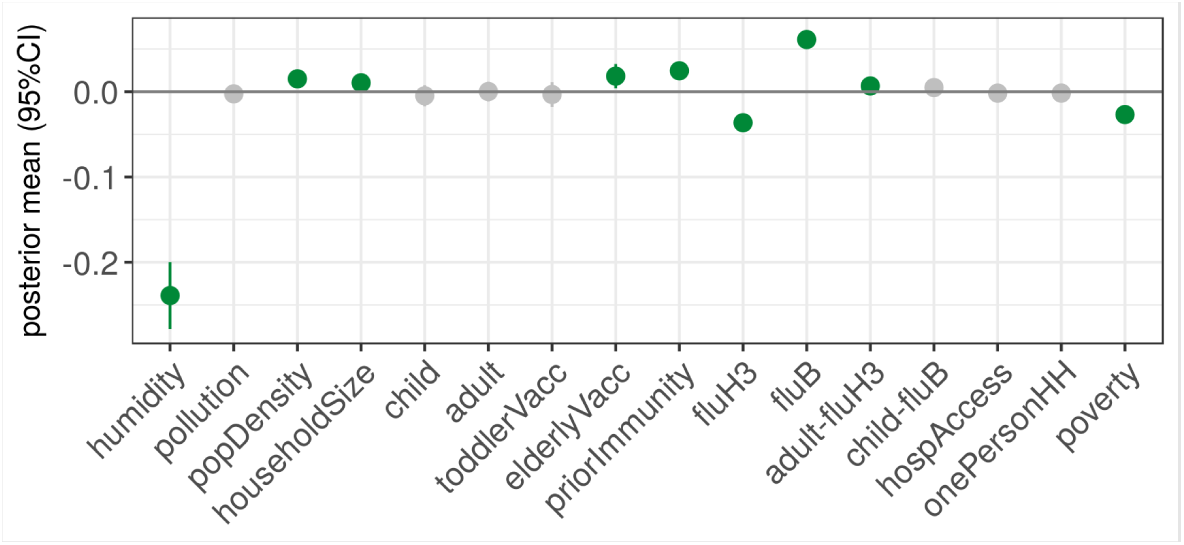
For the total population multi-season epidemic duration models, these are the 95% credible intervals for the posterior distributions of the socio-environmental coefficients and B) measurement-related coefficients.

**Appendix 3 Figure 7.**
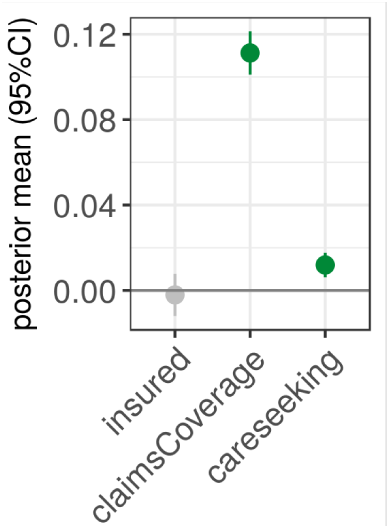
For the total population multi-season epidemic duration models, these are the 95% credible intervals for the posterior distributions of measurement-related coefficients.

## Appendix 4

### Comparison of disease burden metrics

**Appendix 4 Figure 1.**
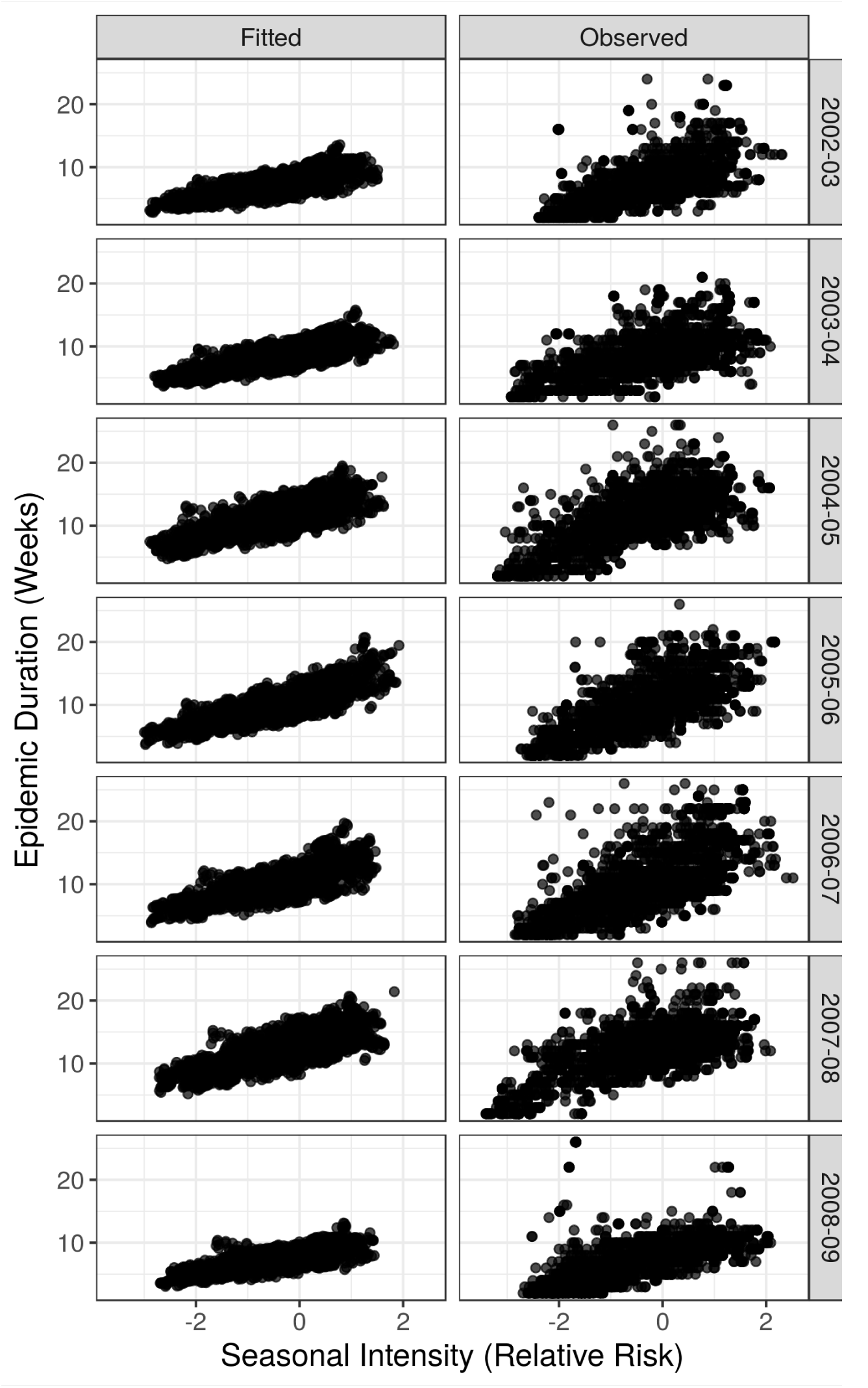
Comparison of epidemic duration and relative risk for seasonal intensity among fitted (left) and observed (right) values.

## Appendix 5

### Model predictors Checks for multicollinearity

We checked for multicollinearity among predictors by examining Spearman rank crosscorrelation coefficients between all pairs of final model predictors (excluding interaction terms). No single pair had a linear correlation coefficient that exceeded a magnitude of 0.6.

**Appendix 5 Figure 1.**
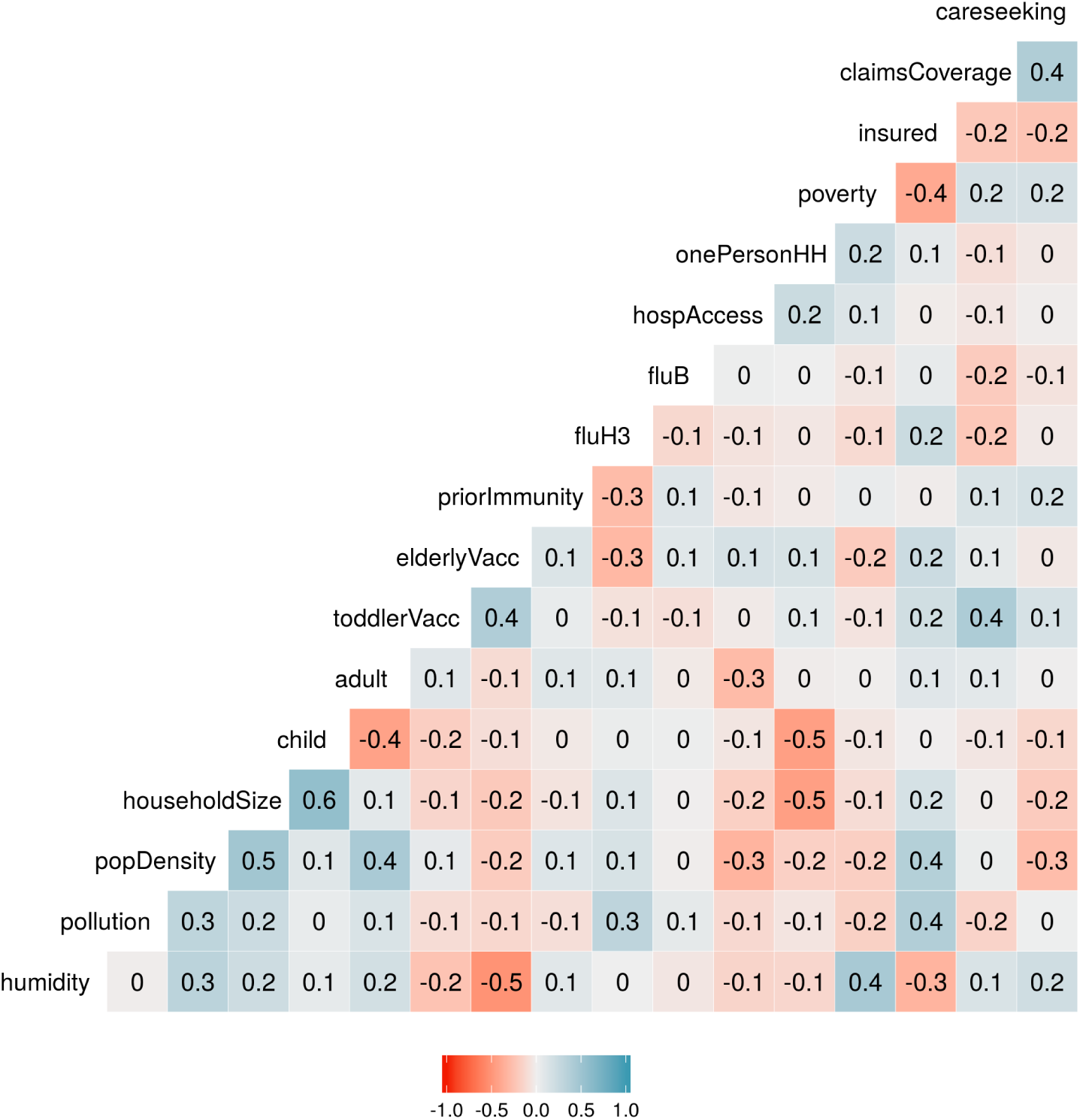
Spearman rank cross-correlation matrix for all pairs of final model predictors.

Additionally, we ran our multi-season seasonal intensity model with each coefficient individually. Multicollinearity between predictors may sometimes be detected when a predictor significantly deviates from zero in the single predictor model, but does not appear to have an effect in a multivariate context. Some predictors (pollution, popDensity, fluB) that were significant in the single predictor context no longer had an effect in our complete model (and vice versa for householdSize and child). Nevertheless, all of these predictors had small effect sizes in both single and multivariate models, and the other predictors that were significant in both models retained effect sizes with the same order of magnitude and directionality.

**Appendix 5 Figure 2.**
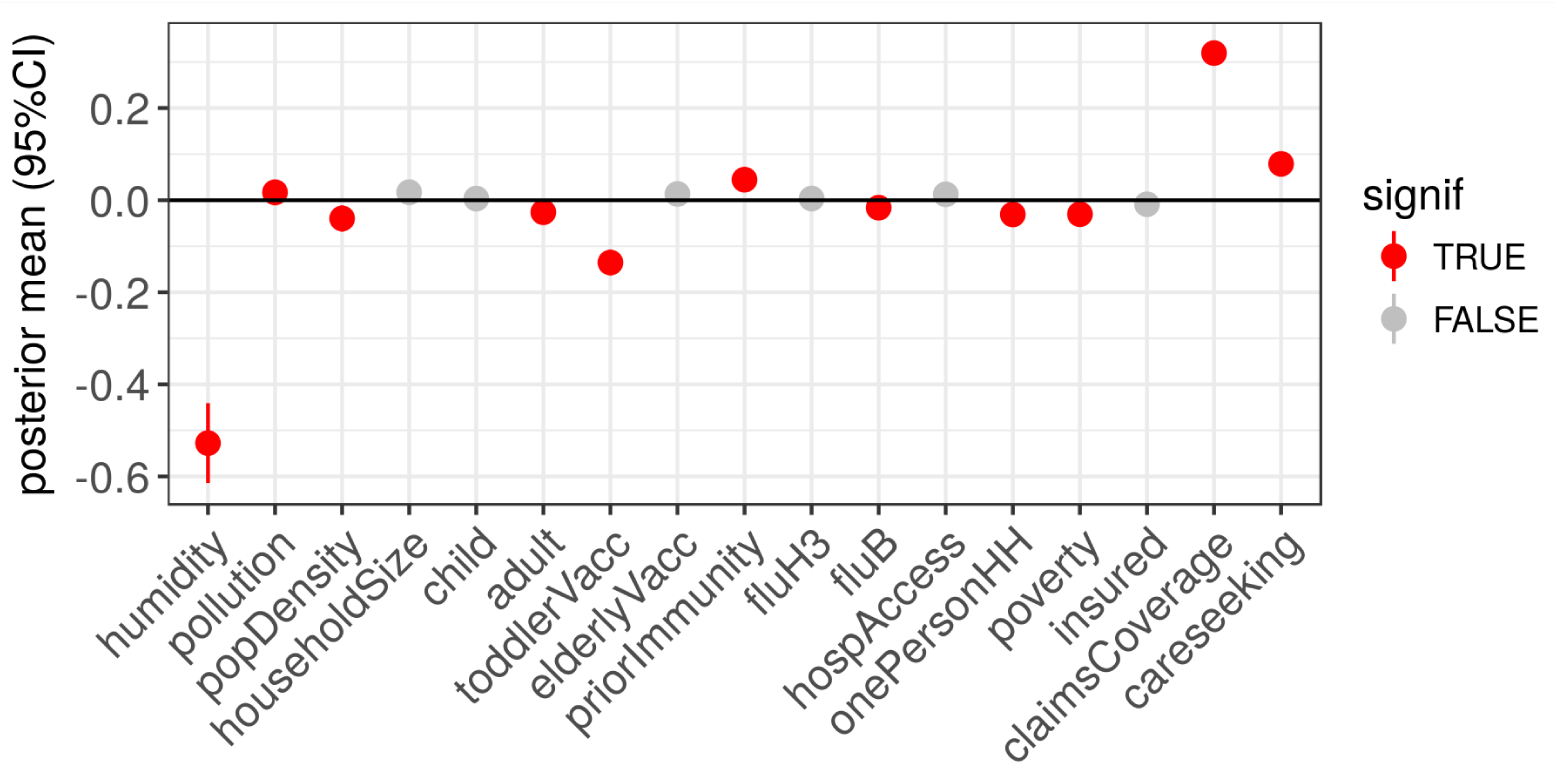
These are the 95% credible intervals among multi-season models with a single predictor for seasonal intensity.

### Medical claims coverage

Medical claims database coverage increased over time across each state.

**Appendix 5 Figure 3.**
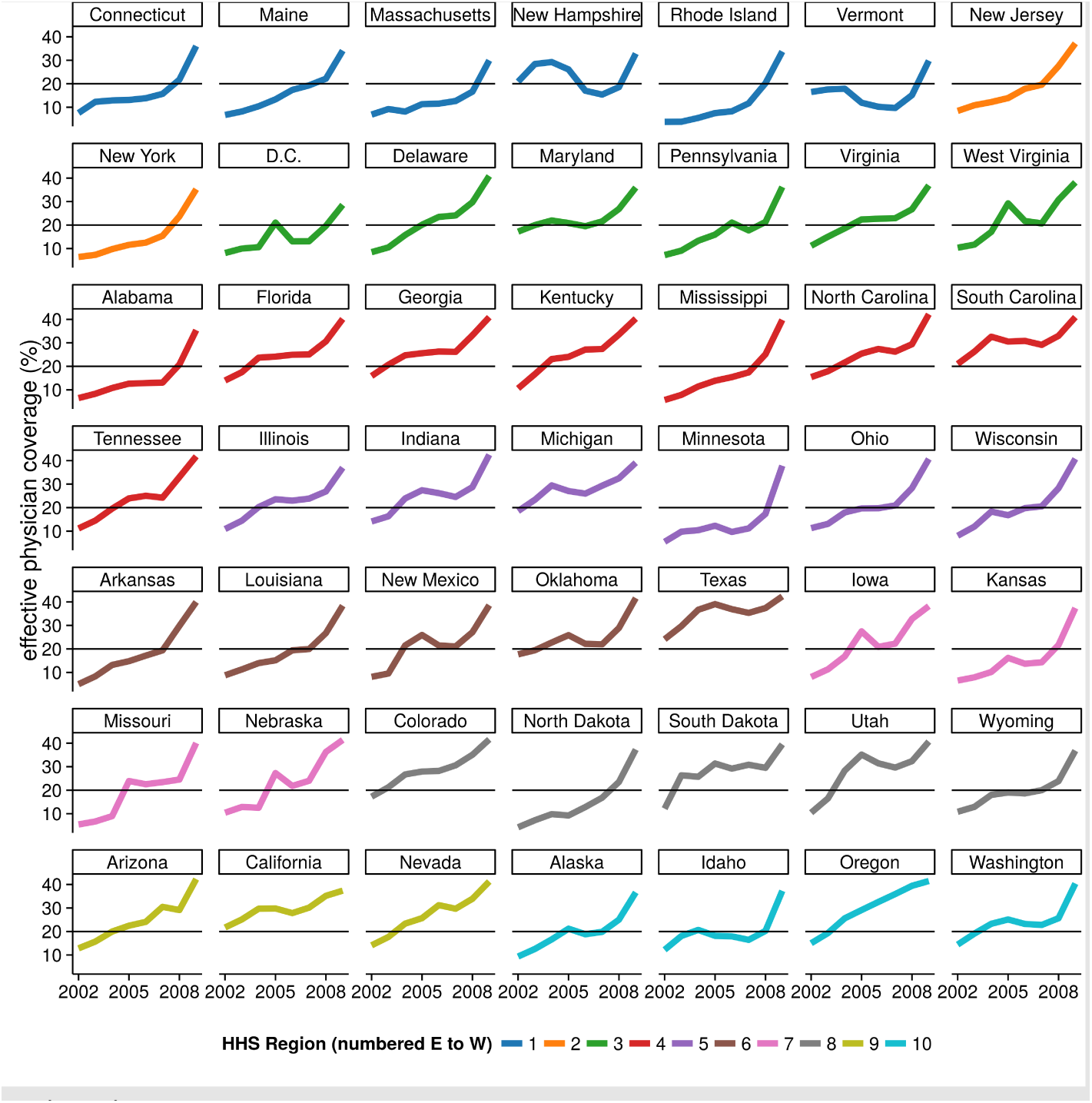
Medical claims database coverage by year and state. Colors represent states that belong to the same HHS region. The black horizontal line at 20% effective physician coverage is a visual guide to ease the comparison of data across panels.

---

**Figure 1-Figure supplement 1.**
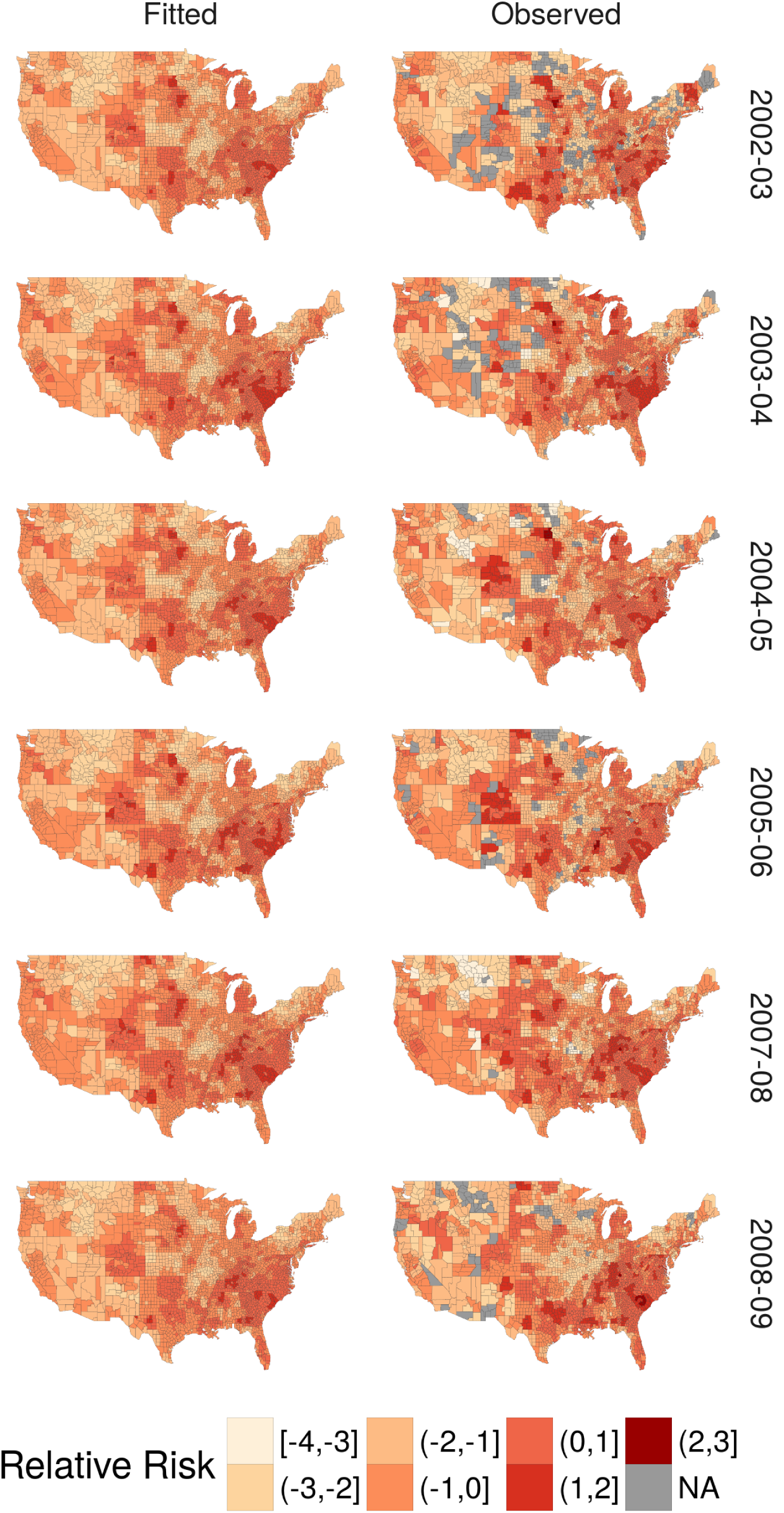
Continental U.S. county maps for fitted (left) and observed (right) relative risk of seasonal intensity for remaining influenza seasons.

**Figure 3-Figure supplement 1.**
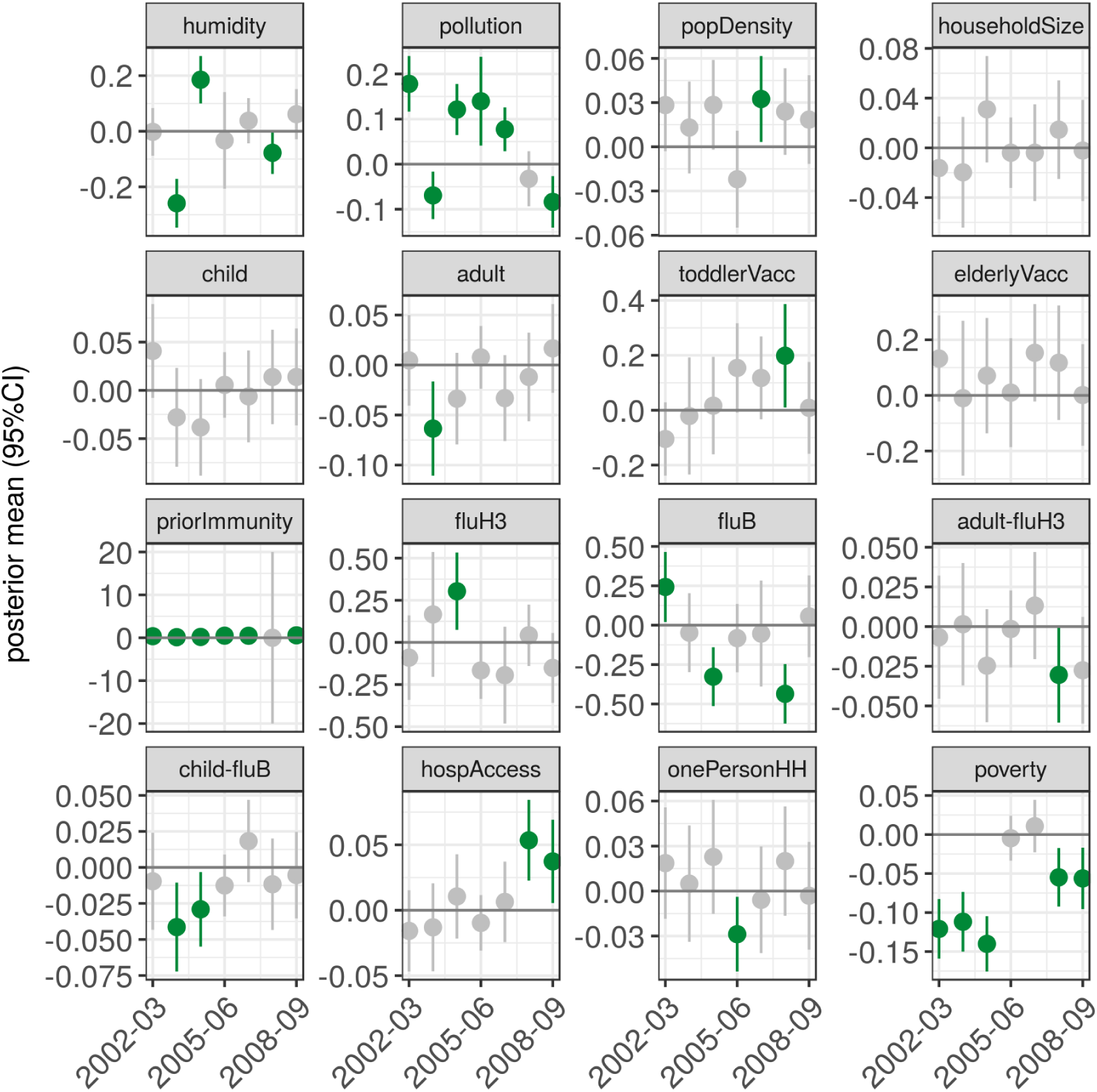
For the total population single-season seasonal intensity models, these are the 95% credible intervals for the posterior distributions of the socioenvironmental coefficients.

**Figure 3-Figure supplement 1.**
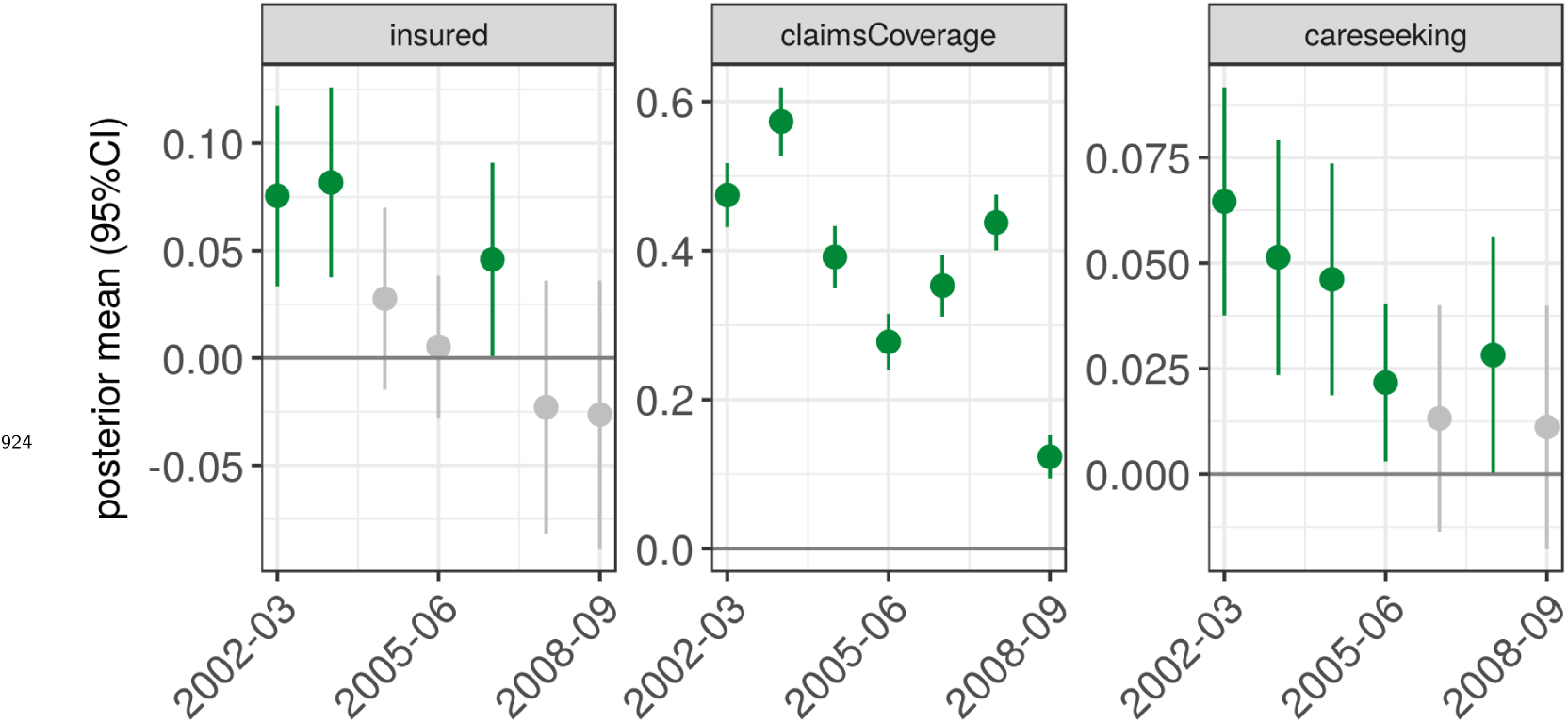
For the total population single-season seasonal intensity models, these are the 95% credible intervals for the posterior distributions of the measurement coefficients.

**Figure 4-Figure supplement 1.**
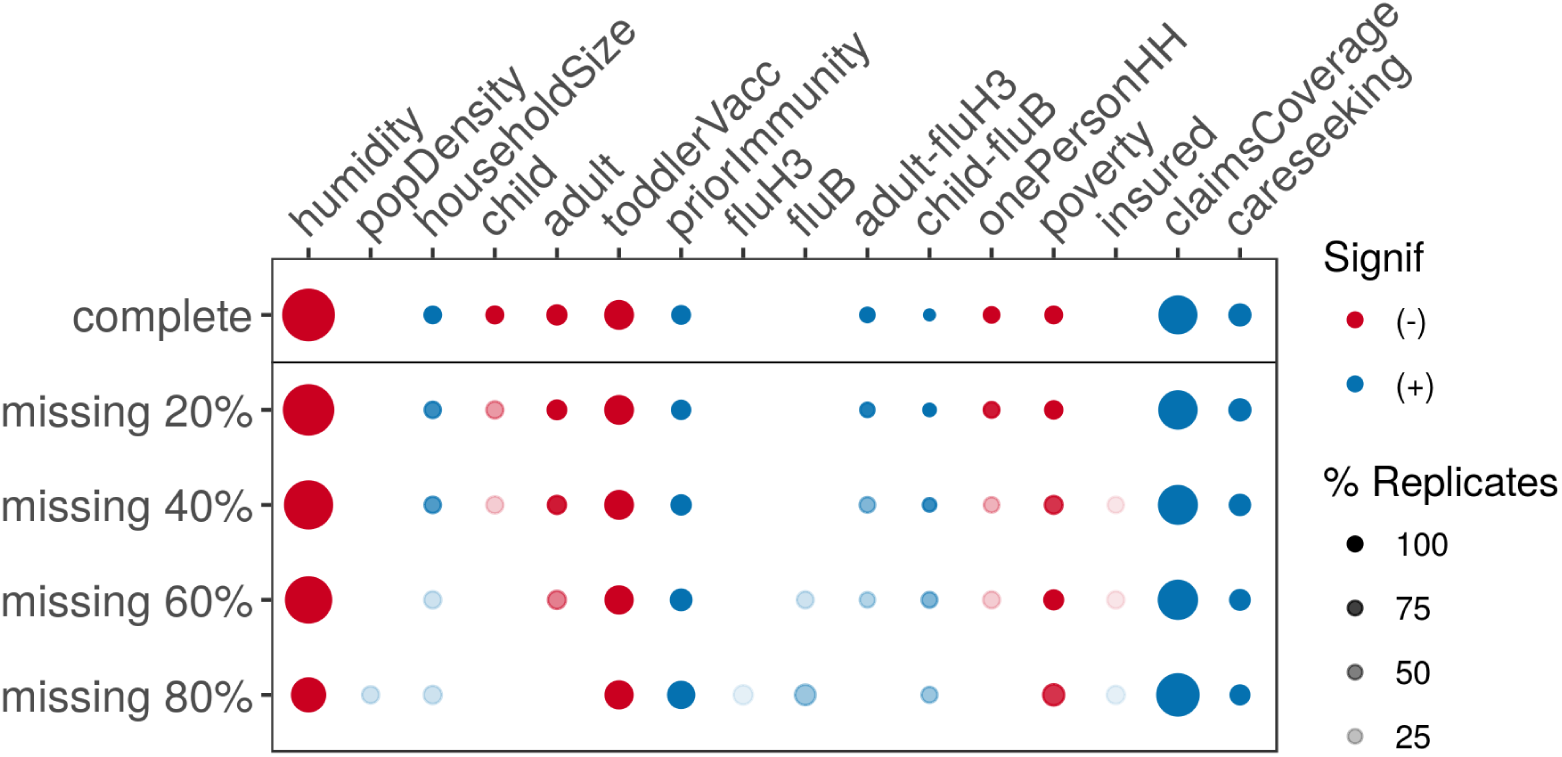
Diagram indicating changes to model inference as fewer moving-location sentinels reported data.

**Figure 4-Figure supplement 2.**
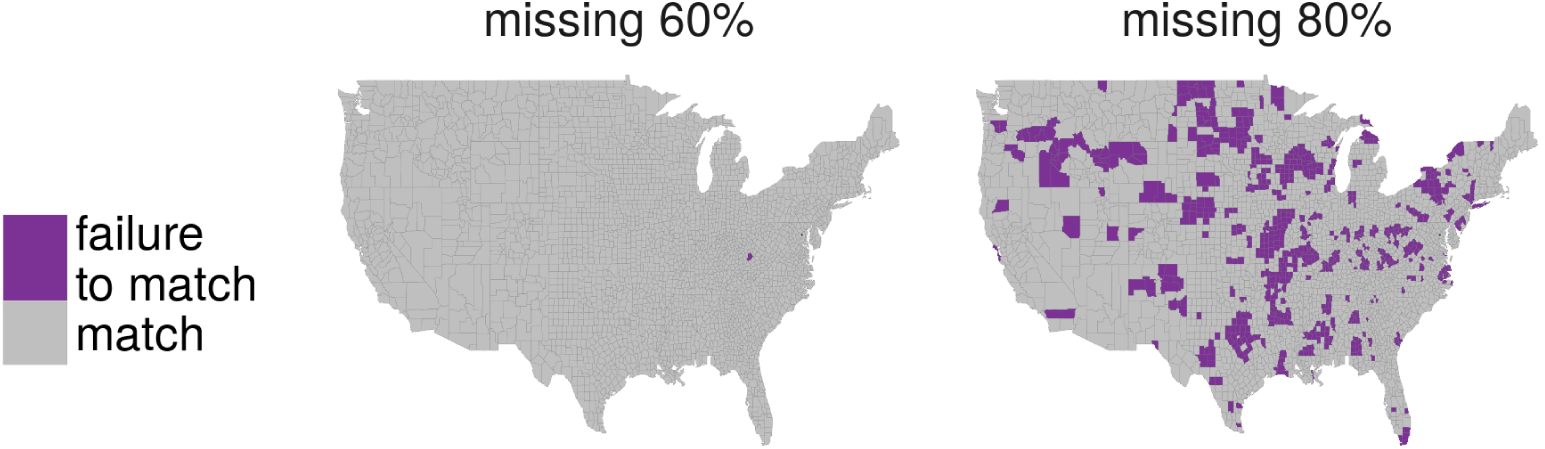
Map of model prediction match between the complete model and the 60% and 80% missing levels for moving-location sentinels.

**Figure 4-Figure supplement 3.**
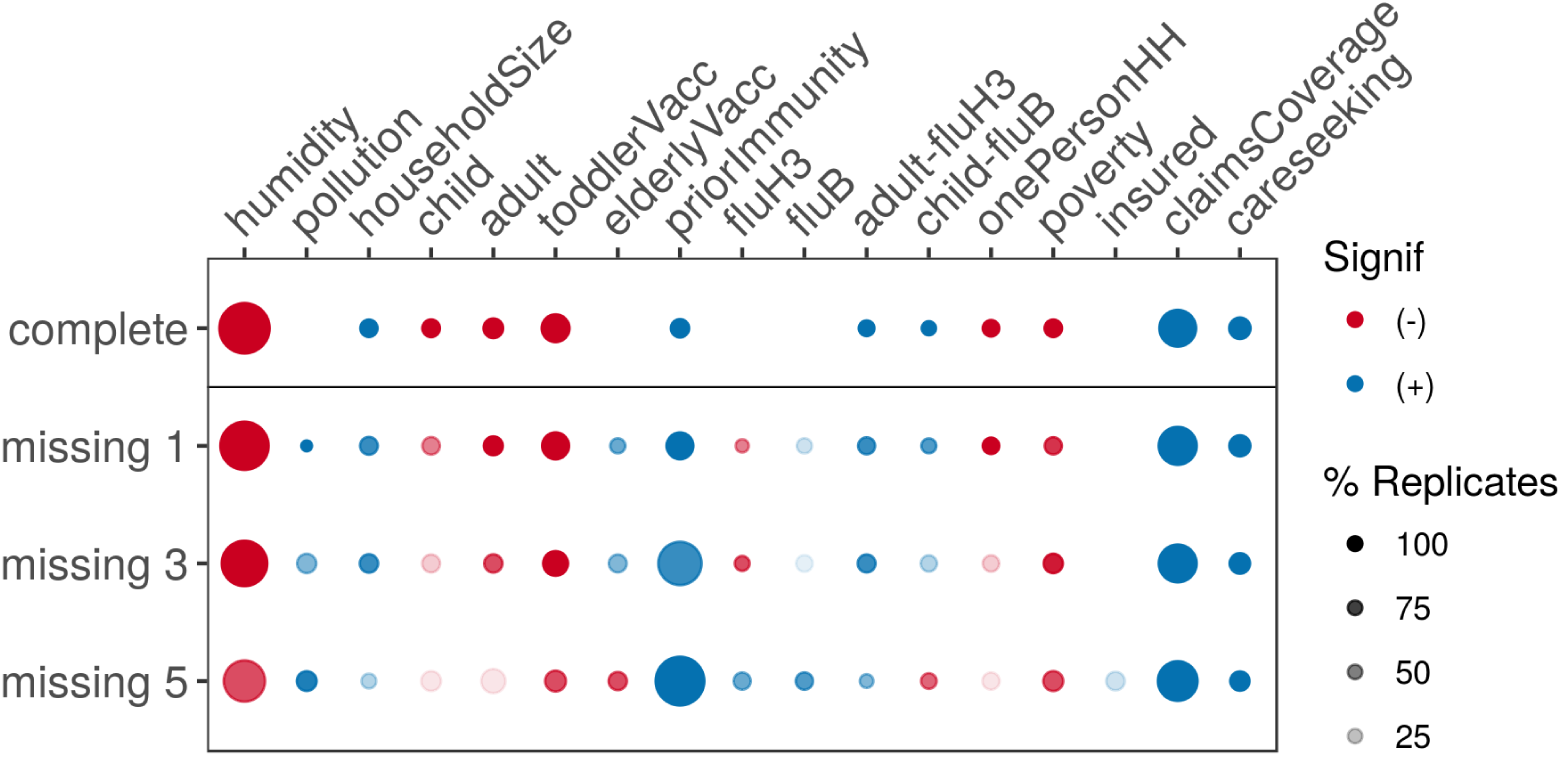
Diagram indicating changes to model inference as historical seasons were randomly removed from the model.

**Figure 4-Figure supplement 4.**
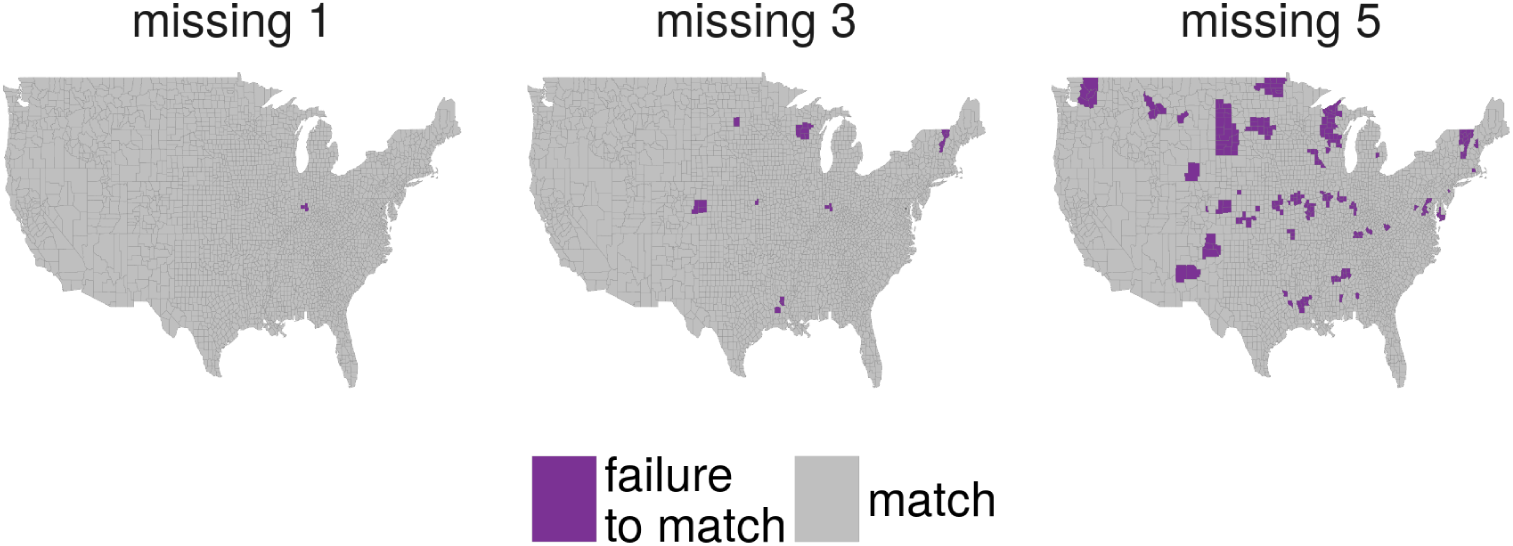
Map of model prediction match between the complete model and models missing one, three, or five historical flu seasons.

